# Cellular Bases of Olfactory Circuit Assembly Revealed by Systematic Time-lapse Imaging

**DOI:** 10.1101/2021.05.04.442682

**Authors:** Tongchao Li, Tian-Ming Fu, Hongjie Li, Qijing Xie, David J. Luginbuhl, Eric Betzig, Liqun Luo

## Abstract

Neural circuit assembly features simultaneous targeting of numerous neuronal processes from constituent neuron types, yet the dynamics is poorly understood. Here, we use the *Drosophila* olfactory circuit to investigate dynamic cellular processes by which olfactory receptor neurons (ORNs) target axons precisely to specific glomeruli in the ipsi- and contralateral antennal lobes. Time-lapse imaging of individual axons from 28 ORN types revealed a rich diversity in extension speed, innervation timing, and ipsilateral branch locations, and identified that ipsilateral targeting occurs via stabilization of transient interstitial branches. Fast imaging using adaptive optics- corrected lattice light-sheet microscopy showed that upon approaching target, many ORN types exhibit “exploring branches” consisted of parallel microtubule-based terminal branches emanating from an F-actin-rich hub. Antennal nerve ablations uncovered essential roles for bilateral interactions in contralateral target selection, and for ORN axons to facilitate dendritic refinement of postsynaptic partner neurons. Altogether, these observations provide cellular bases for wiring specificity establishment.

## INTRODUCTION

The proper function of the nervous system relies on the precise assembly of neuronal circuits. During development, individual neurons extend their axons and dendrites, often across large distances, to match with their synaptic partners. Axons and dendrites are led by growth cones, which navigate through complex extracellular environment at each step of their journey. Numerous neurons are performing this act simultaneously within any given neural region. While great strides have been made in the past decades to identify molecules that control axon guidance, dendrite elaboration, and target selection (reviewed in Jan and Jan, 2010; Kolodkin and Tessier-Lavigne, 2011; Sanes and Zipursky, 2020), the cellular contexts within which most wiring molecules act during the circuit assembly process are not well characterized.

Because of its high temporal resolution, time-lapse imaging has been utilized to define key cellular events in neuronal wiring. Notable examples include discovering growth cone dynamics in tissue culture (Harrison, 1910); identifying guidepost cells for axon guidance in grasshopper limb bud (Bentley and Caudy, 1983); characterizing distinct growth cone dynamics of retinal ganglion cells at the optic chiasm in mice (Godement et al., 1994) or tectal targets in *Xenopus* (Harris et al., 1987); identifying repulsive interactions between sensory dendrites and axons that tile *Drosophila*, zebrafish, and *C. elegans* body surface (Grueber et al., 2003; Sagasti et al., 2005; Smith et al., 2010); investigating the relationship between dendritic growth with synapse formation in the fish and amphibian retinotectal systems (Niell et al., 2004; Haas et al., 2006); and defining the rules and molecular mechanisms of target selection for *Drosophila* photoreceptor axons (Langen et al., 2015; Akin and Zipursky, 2016). Most studies have focused on a single group of cells at a specific developmental stage. It remains unclear how growth cone dynamics of the same neuron change at different stages of circuit assembly, and the extent to which different types of neurons in the same circuit follow the same rules.

The *Drosophila* olfactory system has served as an excellent model for investigating the mechanisms by which neural circuits are assembled. Axons of about 50 types of olfactory receptor neurons (ORNs) and dendrites of 50 types of second-order olfactory projection neurons (PNs) form 1-to- 1 connections in 50 discrete, stereotyped, and individually identifiable glomeruli in the antennal lobe, so as to relay olfactory information from the periphery to the brain (reviewed in Vosshall and Stocker, 2007). During the assembly of the adult olfactory system, PNs first extend their dendrites and establish a coarse map by the positioning of their dendrites (Jefferis et al., 2004). ORN axons then arrive and choose either the dorsolateral or ventromedial trajectory to circumnavigate the antennal lobe, cross the midline, and invade the ipsi- and contralateral antennal lobes to find their synaptic partners (Jefferis et al., 2004; Joo et al., 2013) (Figure 1A). A multitude of cellular and molecular mechanisms have been identified that direct dendrite and axon targeting of selected PN and ORN types (reviewed in Hong and Luo, 2014); some of the molecules and mechanisms first discovered in the fly olfactory system have subsequently been found to be conserved in wiring the mammalian brain (e.g., Hong et al., 2012; Berns et al., 2018). Still, we are far from understanding the developmental algorithms that orchestrate the precise wiring of 50 pairs of ORN and PN types. Time-lapse imaging can provide cellular context in which wiring molecules exert their functions. Moreover, the heterogeneity of neuron types and the ease of identifying them based on their stereotyped glomerular targets enable us to examine the degree to which different neuron types use the same wiring rules.

**Figure 1.**
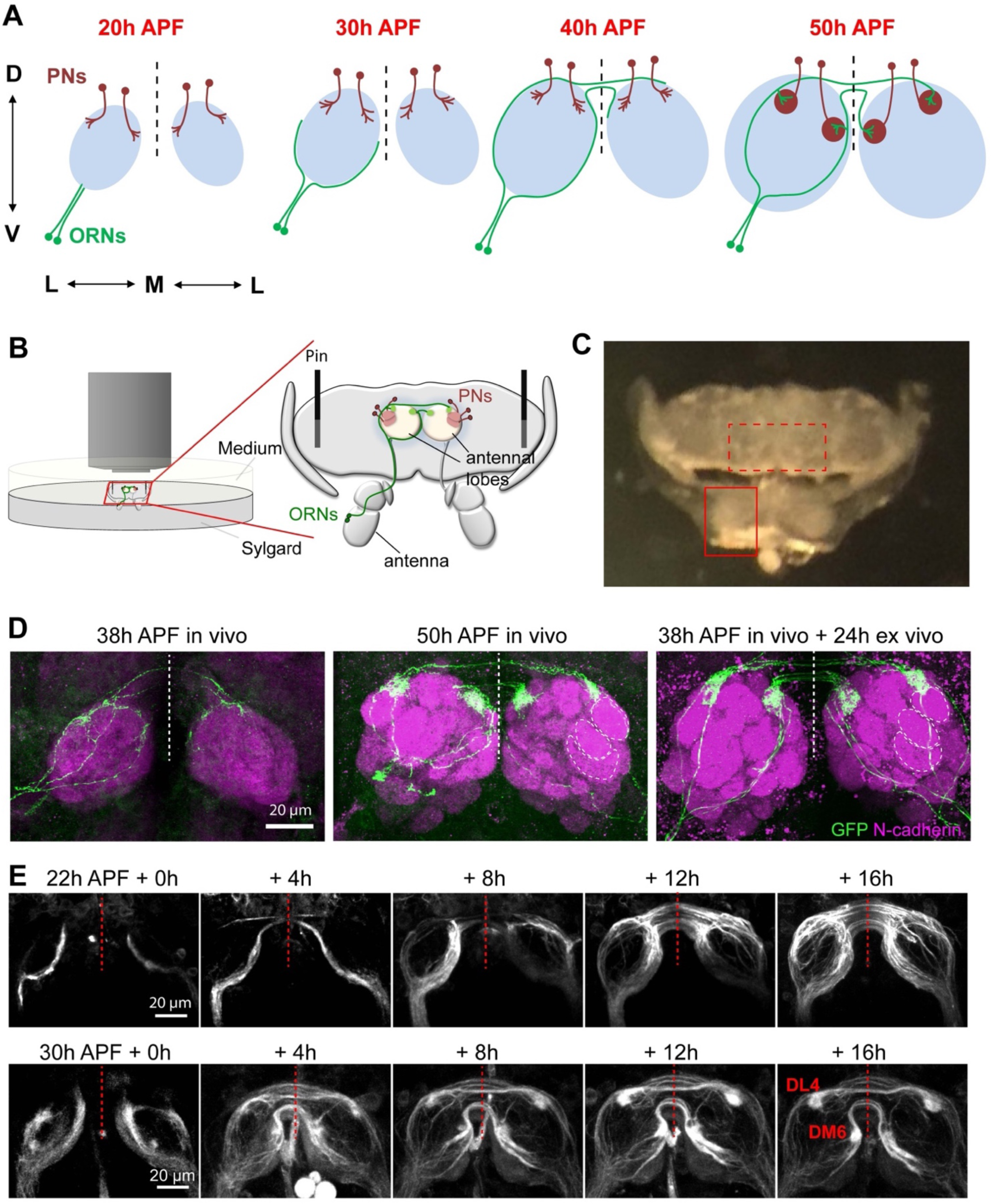
Antennae–brain explants enable live imaging of olfactory circuit assembly. (A) Schematic showing sequences of olfactory circuit assembly in the pupal brain. Abbreviation in this and all subsequent figures: ORN, olfactory receptor neuron; PN, projection neuron; h APF, hours after puparium formation; D, dorsal; V, ventral; M, medial; L, lateral. Dashed vertical lines always indicate the midline of the brain. (B) Schematic showing imaging setting for the antennae–brain explant. (C) A bright-field image shows dissected antennae–brain explant with dorsal side facing upward. Solid and dashed boxes mark the antennal lobes in the brain and one antenna, respectively. (D) Max intensity projections of confocal images from brains dissected at 38h APF (left), 50h APF (middle), and an explant dissected at 38h APF followed by 24h culture ex vivo (right). *AM29-GAL4*+ ORN axons are sparsely labeled by mCD8-GFP using MARCM. Neuropils are visualized using N-cadherin. Dashed circles in the middle and right images mark the boundaries of DA1, VA1d, and VA1v glomeruli in one of the two antennal lobes in each brain. (E) *Images* from *time-lapse* videos taken from a two-photon microscope at the indicated time points of *AM29-GAL4;UAS-mCD8-GFP* explants dissected at 22h APF (top row) and 30h APF (bottom row). The primordial DM6 and DL4 glomeruli are marked. See Figure S1 for related data.

Here, based on a previously established protocol for studying the fly visual system in explant cultures (Ozel et al., 2015), we developed an antennae–brain explant preparation that recapitulates the precision by which the olfactory circuit is assembled in vivo. Time-lapse imaging of the targeting process of 90 single ORN axons from 28 types revealed heterogenous targeting behaviors at multiple steps that contribute to the eventual wiring specificity. <u>A</u>daptive <u>o</u>ptics-lattice light-<u>s</u>heet <u>m</u>icroscopy (AO-LLMS) enabled us to image singly labeled ORN axons with high spatiotemporal resolution and to discover a novel axon terminal structure prior to ORN axons reaching targets. We also found that cytoskeletal organization of ORN axon terminals differs substantially from that of the classic growth cones from neurons in primary culture and varies between different developmental stages. Finally, unilateral and bilateral ablation of ORN axons uncovered essential roles for bilateral interactions in contralateral target selection, and for ORN axons to facilitate dendritic refinement of PNs.

## RESULTS

### An antennae–brain explant system for time-lapse imaging of olfactory circuit assembly

The wiring specificity of the adult *Drosophila* olfactory circuit is established during the first half of the ∼100-hour pupal stage (Jefferis et al., 2004), when the brain is covered with opaque fat bodies inside the pupal case that prevent fluorescence imaging using two-photon microscopy. To obtain high-quality images of the developing olfactory circuit from live tissues, we established an explant system containing the pupal brain, antennae, and their connecting nerves (Figure 1B, C). This was achieved by cutting out the cuticle on the top and bottom sides of the brain using microscissors and removing all fat bodies between the brain and the cuticle. Residual cuticle that connects the optical lobe of the brain and antennae was left to maintain the integrity of the antennal nerves (Figure 1C). The dissected antennae– brain explant was then immobilized on a sylgard plate and cultured for subsequent 24–48 hours.

To assess the degree to which olfactory circuit development ex vivo mimics in vivo conditions, we monitored targeting of single axons from two specific ORN types to the DL4 and DM6 glomeruli labeled by *AM29-GAL4* (Endo et al., 2007) using the MARCM strategy (Lee and Luo, 1999), counterstained with a neuropil marker N-cadherin (Figure 1D). At 38 hours after puparium formation (h APF) in vivo, individual glomeruli had not developed as assessed by the neuropil staining, and ORN axons were still finding their ways to the eventual targets (Figure 1D, left). At 50h APF in vivo, antennal lobes were substantially larger, glomeruli were individually identifiable by the neuropil staining, and ORNs of a given type terminate their axons in one of 50 glomeruli in the ipsi- and contralateral antennal lobes (Figure 1D, middle). When we dissected brain and antennae at 38h APF and cultured the explant for 24h ex vivo, we found that the antennal lobe volumes increased compared to 38h APF in vivo, individual glomeruli were readily identifiable, and *AM29-GAL4*+ ORN axons elaborate their terminals at positions similar to the DL4 and DM6 glomeruli in vivo (Figure 1D, right). *AM29-GAL4*+ ORN axons occasionally targeted to two additional regions outside the DM6 and DL4 glomeruli in culture (Figure S1A, B). Interestingly, minor targeting to similar regions was also observed in vivo during the intermediate developmental stages and became less frequent at later developmental stages (Figure S1A, B). Taken together, these data indicate that the olfactory circuit develops similarly in our explant culture as in vivo, albeit at a slower pace (requiring about 2× the time).

To test whether the explant could be subjected to two-photon microscopy imaging for an extended period of time without affecting the development of the olfactory circuit, we imaged axons of all DM6 and DL4 ORNs using *AM29-GAL4* driven mCD8-GFP reporter expression (Figure 1E, Movie 1), or axons of all ORN types using the *pebbled-GAL4* driver (Figure S1C). The growth and targeting of ORN axons were not disrupted by continuous imaging for 24h at a rate of once every 20 min, suggesting little photodamage at this imaging frequency. Thus, this explant system can be used for long-term live imaging study of olfactory circuit assembly.

### Imaging glomerular targeting of individual ORNs

We next attempted to perform high-resolution live imaging of individual axons of multiple ORN types. Each ORN type comprises on average ∼30 ORNs from each antenna (∼60 on both sides), making it impossible to distinguish single ORN terminals even with a GAL4 line that only labels one ORN type. Although MARCM (Lee and Luo, 1999) allows sparse labeling of single ORN axons (Figure 1D), it could not produce strongly labeled individual axons at early stages of axon growth, likely due to the perdurance of GAL80 during the limited time between the last cell division of the progenitor cells and ORN axon targeting. To achieve sparse and strong labeling from early development, we modified the FLPout strategy (Wong et al., 2002) by using *FRT* sites with reduced recombination efficiency (Figure 2A). By introducing different point mutations of *FRT* (Senecoff et al., 1988), we generated two reporters, *UAS-FRT^10^-stop-FRT^10^-mCD8-GFP* (Figure S1D) and *UAS-FRT^100^-stop-FRT^100^-mCD8-GFP* (Figure 2B) that were about 10 and 100 times less efficient than the wild type FLP/*FRT* system, respectively.

**Figure 2.**
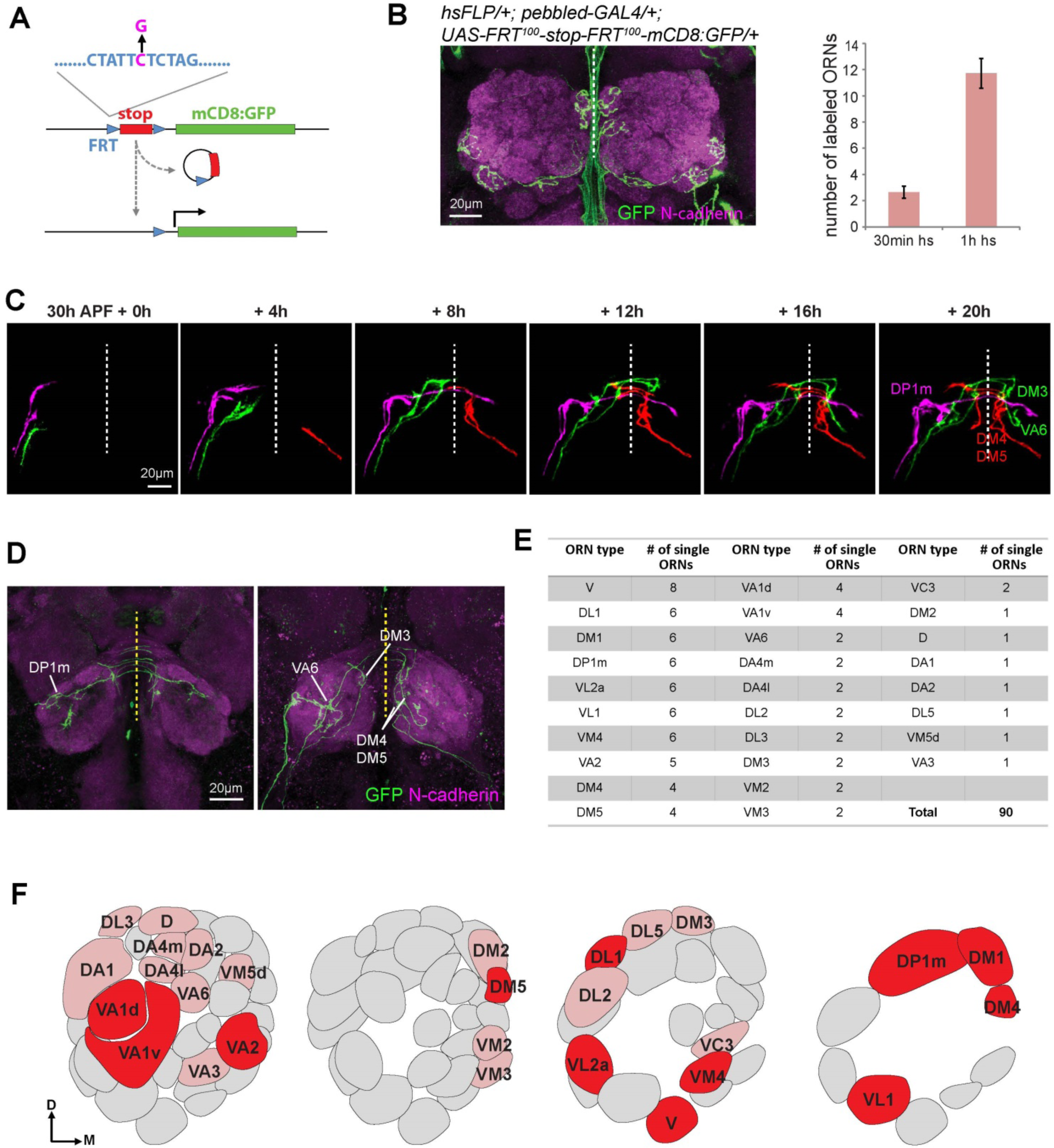
Live imaging glomerular targeting of single axons of multiple ORN types. (A) Schematic showing strategy of sparse labeling using FLPout with mutant *FRT* sites with reduced recombination efficiency. The C→G mutation (magenta) in the *FRT* site reduces recombination efficiency by ∼100 fold. (B) Max intensity projections of confocal images showing four ORNs labeled in *hsFLP,pebbled- GAL4;UAS-FRT^100^-stop-FRT^100^-mCD8- GFP/+* adult brain after heat shock for 30 min at 0h APF (left). The number of labeled ORNs are quantified by different times of heat-shock (hs, right). Green signal in the midline region is from non-specific staining at the surface of the brain. (C) Example *images* from *time-lapse* videos taken from a two-photon microscope at the indicated time points of sparsely labeled ORN axons in *pebbled-GAL4,hsFLP;UAS-FRT^100^-stop-FRT^100^-mCD8-GFP* explant. Axons from different ORNs were followed through time-lapse videos and pseudo-colored manually. (D) Max intensity projections of confocal images of the posterior and anterior antennal lobes from the fixed explant shown in (C) following 24h culture. The genetic identity of each axon was identified by its glomerular target. (E) A summary table for single ORNs live-imaged for each type, with the number of imaged single ORNs indicated. (F) An antennal lobe map with glomeruli corresponding to imaged ORNs labeled. Dark and light red glomeruli represent ORN types with >3 and <3 ORNs imaged, respectively. See Figures S2 and S3 for related data.

Using *UAS-FRT^100^-stop-FRT^100^-mCD8-GFP*, we could randomly label a few ORNs out of ∼1500 per antenna expressing mCD8-GFP driven by the pan-ORN *pebbled-GAL4* driver by heat shock-induced FLP expression from the *hsFLP* transgene (Figure 2B). From ∼1000 pupal brain dissected at 30h APF, we selected 75 brains in which one or few axons that can be individually separated just arrived at the antennal lobe, and performed time-lapse imaging in explant culture at the frequency of once every 20 min for the next 24h, which covered the entire targeting process of each ORN axon in the antennal lobe (Figure 2C). The identity of each ORN was determined by subsequent immunostaining of the fixed explants following the 24h culture, counterstained with the neuropil marker N-cadherin (Figure 2D). We extracted 90 single ORNs from these 75 brains, covering 28 ORN types (Figure 2E, F; Figure S2; Movie 2). These time-lapse images provided a valuable dataset to analyze how individual ORN axons find their target regions, as detailed in the next four sections.

### Axons of different ORN types reach the antennal lobe following a temporal sequence

Previous studies indicate that pioneering ORN axons reach the antennal lobe around 18h APF, and by 48h APF, all ORN axons complete their glomerular targeting (Jefferis et al., 2004), but it is unknown whether there is any heterogeneity in the timing of axon development for different ORN types. The 90 single ORNs we selected on the basis of axon arrival at the antennal lobe at 30h APF covered only a subset of ORN types at different frequencies (Figure 2E, F), suggesting heterogeneity in the timing of axon arrival at the antennal lobe for different ORN types.

To extend this finding, we compared explant cultures initiated at 26h, 30h, and 34h APF, and determined the glomerular identity after 24–48h culture for those axons that just reached the antennal lobe at 26h or 30h APF, or those that reached the antennal lobe after 34h APF (Figure S3A). This allowed us to determine the glomerular identity for axons that arrive early (at 26h APF), middle (at 30h APF) or late (after 34h APF). We found that early-arriving ORN axons tended to target more posterior glomeruli, while late-arriving ORN axons tended to target more anterior glomeruli (Figure S3B, C). Thus, different ORN types not only target their axons to spatially segregated glomeruli, but their axons also follow a temporal sequence in their arrival at the antennal lobe. This temporal segregation should in principle reduce the complexity of cellular context with which axons of each ORN type interact as they navigate. We note that posterior-targeting ORN types have more distinct transcriptomes at 24–30h APF compared with other ORN types (McLaughlin et al., 2021), suggesting genetic control for the timing of axon development.

### ORN axon growth slows down at specific choice points

Previous live imaging studies have revealed that axons reduce their extension speed at specific choice points. For example, vertebrate retinal ganglion cell axons slow their extension at the optic chiasm, a choice point along their trajectory (Godement et al., 1994; Hutson and Chien, 2002; Bak and Fraser, 2003). Do ORN axons change their speed at specific points during their journey? Are there type-specific differences? To address these questions, we measured the axon growth speed of 5 ORN types that had more than 3 samples, targeting three lateral glomeruli (DL1, DP1m and VL2a; green shades in Figure 3A, B) and two medial glomeruli (DM1 and DM4; red and magenta in Figure 3A, B). To analyze ORN axon growth in the ipsilateral antennal lobe, we defined the last imaging scan when axons have not crossed the midline as time 0, and quantified the change of axon length in each imaging scan interval relative to time 0. We found that ORN axons grew at an average speed of 2–6 µm/20 min imaging interval (0.1–0.3 µm/min) when they circumnavigated the ipsilateral antennal lobe (Figure 3B, left). This speed reduced to 0–2 µm/20 min right before midline crossing for all 5 types (solid arrow in Figure 3A, B), suggesting that ORN axons reduce their growth speed before midline crossing in general.

**Figure 3.**
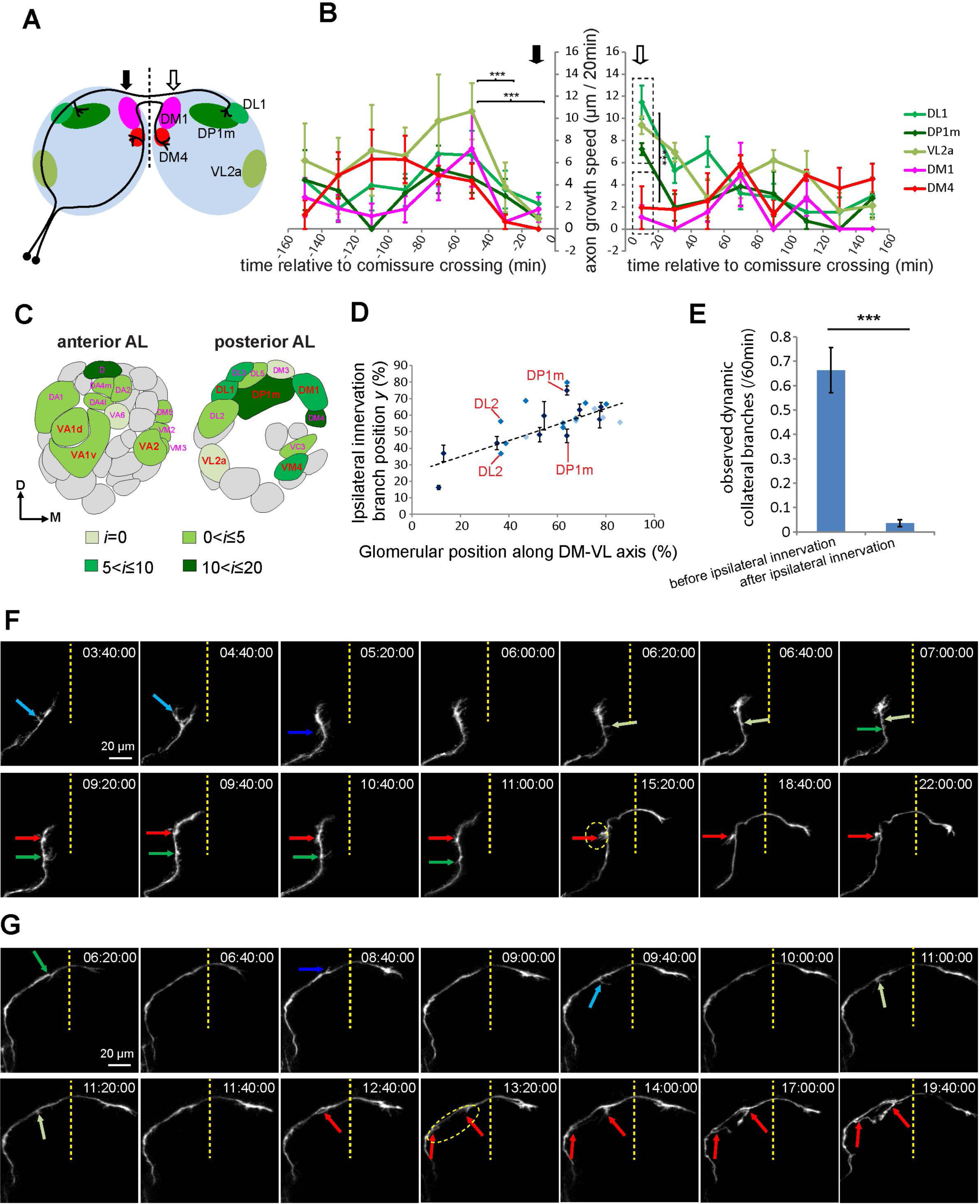
Quantitative analysis of axon targeting events from time-lapse imaging of single ORN axons. (A) Antennal lobe map showing the DL1, DP1m, VL2a, DM1, and DM4 glomeruli in different colors. Solid and open arrows mark regions before and after the midline crossing, respectively. (B) The axon growth rate of the 5 ORN types in (A) relative to time of midline crossing. Asterisks in the left chart show statistical significance of the growth speed from all types between corresponding time windows. Asterisks in the right chart show statistical significance between the growth rate of DL1/DP1m/VL2a axons and DM1/DM4 axons within the same time window (0–20 min). Statistics between different time points across all ORN types before midline crossing (left chart) was performed using two-way ANOVA. Statistics between lateral- and medial-targeting ORN axons at the first time point post midline crossing (between dashed-line boxes in right chart) was performed using t-test. ***: P<0.001. Solid open arrows mark time points before and after midline crossing, respectively. n = 5 and 4 for DL1 ORNs and all other ORN types, respectively. (C) The distribution of ORN types with different average *i* value (delayed time for ipsilateral innervation) on a glomerular map, quantified from 90 single ORNs imaged. D, dorsal; M, medial. The types of ORN with n ≥ 3 single ORNs quantified are shown in red, bold, and larger font. The types of ORN n < 3 single ORNs are shown in magenta, regular, and smaller front. (D) Scatter plot showing correlation between the position of centroid of each glomerulus along the dorsomedial (DM)–ventrolateral (VL) axis of the adult antennal lobe and the ipsilateral innervation branch position *y* (see Figure S5A). The dark, medium and light blue dots indicate ORN types with ≥ 3, 2, or 1 sample in our collection, respectively. DL2 and DP1m have two ipsilateral innervation branch points and are indicated in the plot. (E) Quantification of observed dynamic collateral branches (not including the ipsilateral innervating branches) before and after ipsilateral innervation quantified from two-photon microscope based time-lapse imaging of single ORNs (Figure 2E). The following ORN axons were not included in the quantification: VA1d and VA1v axons due to bundling together during targeting; VM2 and VM3 axons due to bundling together during targeting; VL2a, VL1 and VM4 axons due to innervation very close to the entry point in the ipsilateral antennal lobe. t-test; ***, P<0.001. (F, G) Two-photon microscopy-based time-lapse images of a DA2 (F) and a DP1m (G) axon in the indicated time points. Arrows in green and blue colors denote different transient collateral branches. The red arrows mark branches that innervate glomeruli at the end. Dashed line circles outline the future glomeruli. See Figures S4 and S5 for related data.

To analyze ORN axon growth in the contralateral antennal lobe, we defined the time of first imaging scan when the axons left the midline area as time 0. After crossing the midline (open arrow in Figure 3A, B), axon growth speeds differed depending on their locations of their targets. ORN axons targeting the lateral antennal lobe (DL1, DP1m, VL2a), which do not need to make a turn after crossing the midline, grew faster (∼10 µm/20 min) during the first imaging interval after crossing the midline. By contrast, ORNs targeting their axons to the medial antennal lobe (DM1, DM4), which requires a ∼90° turn ventrally, grew slower during the first imaging interval after crossing the midline (Figure 3B, right). Thus, ORN axon growth slows down prior to midline crossing and large turns in their trajectory.

### ORNs innervate ipsilateral glomeruli predominantly via interstitial branching

Selective axon branching enables the same neuron to innervate multiple targets. Axon branching can occur via two distinct cell biological mechanisms: growth cone splitting—the two branches are generated at the same time (Figure S4A, left) or interstitial branching—the second branch (a collateral) extends from an existing axon trunk (Figure S4A, right). Axon branching occurs predominantly via interstitial branching in vertebrate CNS neurons, but axons of dorsal root ganglia sensory neurons branch by growth cone splitting (reviewed in Kalil and Dent, 2014). Since axons of most ORN types innervate the same glomerulus in both the ipsi- and contralateral antennal lobe (Lin et al., 2018), each bilaterally targeting ORN must send one branch (or more, see below) that innervates the ipsilateral glomerulus (referred as ipsilateral branch hereafter) and another branch that extends across the midline to innervate the contralateral glomerulus (Figure 1A).

To determine the cellular mechanism of ORN axon branching, we analyzed from our time-lapse series the interval between when the axon reached its eventual ipsilateral branch point and the first appearance of the ipsilateral branch (Figure S4B). If branches form by growth cone splitting, we would expect the intervals between these two imaging sessions (*i*) to be 0; that is, as soon as the contralateral projecting axon passes the eventual ipsilateral branch point, we should observe the ipsilateral branch already. However, from the 90 single ORNs analyzed, most form the ipsilateral branch with *i* ≥ 1 session of 20 min (Figure 3C). Thus, these ORN axons must innervate their ipsilateral glomeruli via interstitial branching.

We did observe that ORN axons targeting to VL2a (6 single ORN axons), DM3 and VA6 (2 single ORN axons each) glomeruli had *i* = 0 (Figure 3C, S4B). We therefore could not distinguish whether interstitial branching occurred within a 20-min imaging session, or branching occurred via growth cone splitting for these types of ORNs. We also note that different ORN types had considerably different *i* values (Figure 3C). ORNs targeting their axons to ventral glomeruli tended to have smaller *i*, meaning that there was a large time lag between ipsilateral and contralateral glomerular innervation (Figure S4C). ORN types targeting axons to most dorsal glomeruli tended to have large *i*, resulting in nearly simultaneous innervation of ipsilateral and contralateral glomeruli (Figure S4D).

### ORN axons form ipsilateral branches by stabilizing dynamic interstitial branches close to their eventual glomerular targets

Determining the spatiotemporal dynamics of the ipsilateral branch formation can suggest mechanisms of target selection. A priori, several mechanisms can be envisioned. First, the branch point of a given ORN type may be completely genetically specified to be located closest to the final glomerular target, such that the ipsilateral branch has the shortest distance to travel before reaching its target (Figure S5A1). Second, the branch point may be random, and the ipsilateral branch explores a large target region within the antennal lobe before reaching its target (Figure S5A2). Third, ORN axons may produce transient interstitial branches at multiple locations and stabilize the branches that connect with the target (Figure S5A3). To distinguish between these possibilities, we first measured the relative position of the ipsilateral branch point with respect to the entire ipsilateral trajectory from 0 (representing the antennal lobe entry point ventrolaterally) to 100 (representing the midline dorsomedially) using ipsilateral branch point index *y* (Figure S5B). We measured each axon’s curve length in the ipsilateral antennal lobe at the first imaging section after the axon crossed the midline and sent out the ipsilateral branch.

By the time ORN axons crossed the midline, most ORN axons formed ipsilateral branch(es) at one location, except DP1m and DL2 ORNs, which formed ipsilateral branches at two distinct locations and these branches converged onto the same (relatively large) glomerulus (Figure S5C, D). In some ORN types, the ipsilateral branch extended from a single point on the main axon trunk (Figure S5E), while in other ORN types the ipsilateral branch appeared in a continuous area along the axon (Figure S5F). In the latter situation, we used the mid-point of the ipsilateral branching area to determine the ipsilateral branch point index *y*. Comparing *y* of each ORN type and their glomerular location in the adult antennal lobe revealed a coarse trend: ORNs targeting axons to increasingly dorsomedial glomeruli had increasingly larger *y* (Figure 3D, S5D), suggesting that ORN axons form ipsilateral branches close to their future glomerular targets.

Closer examinations of our time-lapse movies revealed that all ORN axons exhibited multiple transient interstitial branches in the ipsilateral antennal lobe before the innervation of ipsilateral glomeruli (Figure 3E). These dynamic interstitial branches were much less frequently observed after the ipsilateral branch reached the target glomeruli (Figure 3E). For ORN types that formed the ipsilateral branch shortly after the main axon passed the ipsilateral branching point (small *i* in Figure 3C), dynamic collateral branching disappeared soon after the ipsilateral branch reached the ipsilateral glomeruli (Figure 3F). For ORN types that had delayed ipsilateral branch formation (large *i* in Figure 3C), dynamic interstitial branching was still observed well after the contralateral branch crossed the midline (Movie 3, Figure 3G). These observations suggest that ipsilateral branches form by stabilizing dynamic interstitial branching along the axons by signals from the target, which in turn inhibits further interstitial branching. This mechanism explains the correlation of positions of branching points and glomeruli described above (Figure 3D).

In addition to the dynamic collaterals, we also observed highly dynamic growth of ipsilateral branches toward the final targeting regions. We extracted the ipsilateral branchs across time from three individual DL1 and DM1 ORNs (Figure S5E, F). While individual neurons of a given ORN type exhibited certain stereotypy—DL1 axons extended ventromedially towards their target, whereas DM1 axons hugged around the branching points because of the proximity of the target to the main ORN axon trajectory—they also exhibited considerable variation in their detailed branching patterns across time. Both DL1 and DM1 ipsilateral branches exhibited dynamic extension and retraction (arrows in Figure S5E, F) instead of steady growth towards the target region. These dynamic changes of the branch terminals suggest that they were actively exploring the local region for target selections before stabilization in the correct targeting region.

### AO-LLSM imaging reveals “exploring branches” of ORN axons before glomerular innervation

The scanning speed and spatial resolution of two-photon microscopy-based imaging limited our ability to examine rapid dynamics of subcellular structures such as the growth cone. We next utilized <u>a</u>daptive <u>o</u>ptics corrected lattice light-<u>s</u>heet <u>m</u>icroscopy (AO-LLSM) that can adaptively correct for optical aberrations caused by the live brain tissues and enables noninvasive volumetric imaging at higher spatiotemporal resolution (Chen et al., 2014; Wang et al., 2014; Liu et al., 2018). To better label the fine structure of single ORN terminals and to increase photostability of the fluorophore, we expressed membrane targeting Halo-tag (Kohl et al., 2014; Sutcliffe et al., 2017) in *peb-GAL4*-based MARCM clones followed by incubation with Janelia Fluor 646 Halo-tag Ligand (Grimm et al., 2017) in explant culture media (STAR Methods). To both image fast dynamics and different stages across long-term development, we imaged every 30 seconds for 14.5 minutes (Movie 4) followed by a break of 1 hour 45.5 minutes, and repeated this image procedure for 24 hours (Figure 4A). The explant was then cultured for another 24 hours before fixing and counter-staining with N-cadherin to reveal the glomerular type of labeled single ORNs. The adaptive optics correction improved the spatial resolution of individual axon branches (Figure 4B), and in combination with fast imaging rate allowed us to capture rapid changes of the fine structure in axon terminals (arrows and arrowheads in Figure 4C).

**Figure 4.**
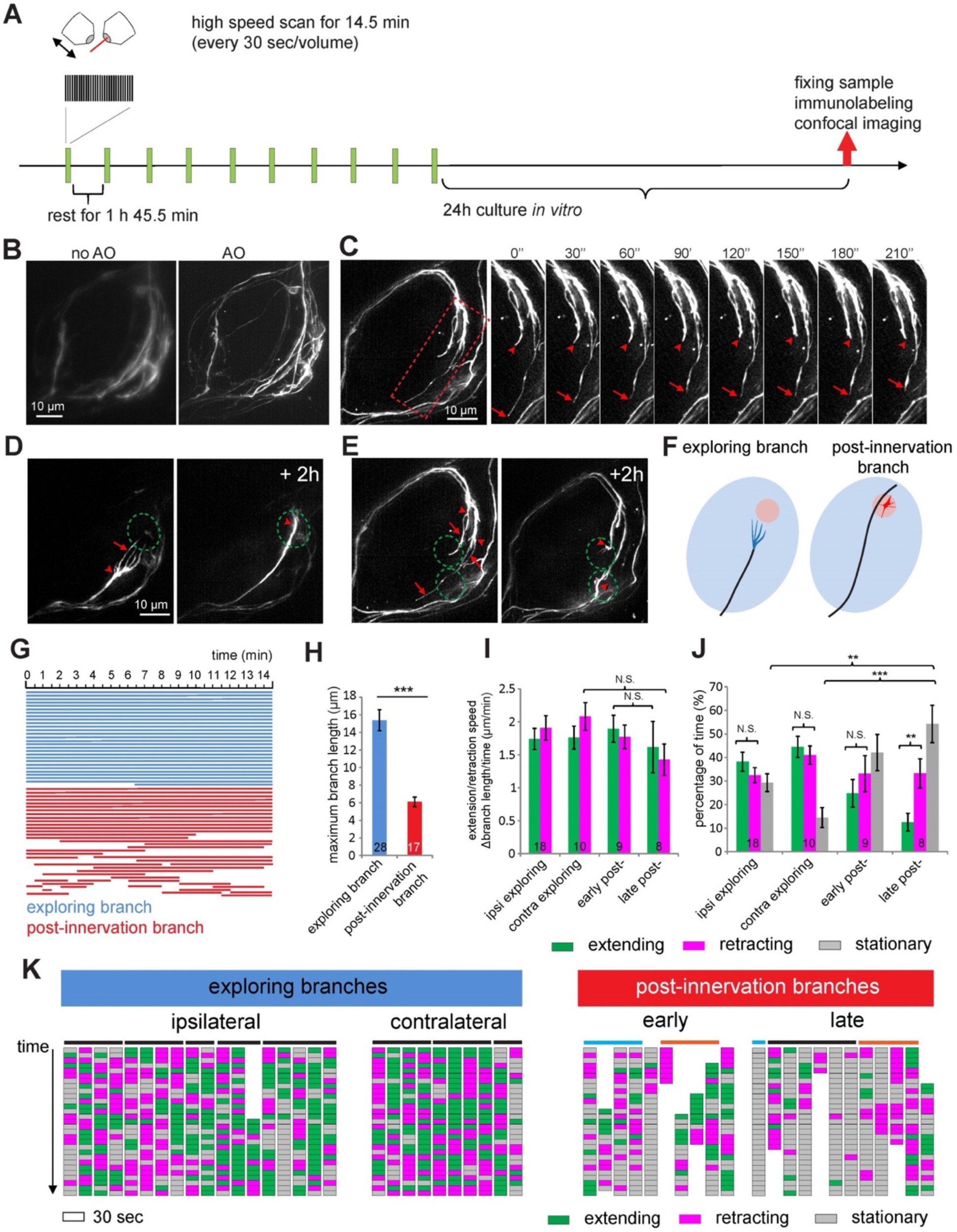
AO-LLSM imaging reveals a novel exploring branch structure of ORN axons. (A) Schematic showing the imaging procedure for AO-LLSM imaging. (B) Two max intensity projections of the same imaged volume of an antennal lobe taken by LLSM without and with adaptive optics (AO) correction. ORNs are sparsely labeled by *pebbled-GAL4* labeled MARCM clones expressing membrane targeting Halo-tag. The explants were dissected at 30h APF, followed by 1h incubation with JF-Halo-646 in culture before imaging. All images in this Figure show explants similarly labeled. (C) Max intensity projection images taken with AO-LLSM every 30 seconds. Time series are from the rectangular box in the still image on the left. Arrows and arrowheads denote the ends of two terminal branches. (D, E) Max intensity projection images taken at two time points with 2h apart from two antennal lobes. Arrows indicate exploring branches and arrowheads mark the hubs where terminal branches converge. Dashed circles denote the future glomeruli. (E) shows the same antennal lobe as (C), with two target glomeruli for two ORN axons. (F) Summary drawing to illustrate exploring branches (blue) and post-innervation branches (red). (G) Lifetime of exploring branches (blue) or post-innervation branches (red). Each line indicates a single branch. The region on the time axis covered by each bar indicates the time window when the branch exists within a 14.5-min fast scanning session. (H) Quantification of the maximal length of the exploring branches or post-innervation branches during the 14.5-min fast scanning sessions. Numbers in the columns indicate numbers of branches quantified (same in I, J). t-test; ***, P<0.001. (I, J) The extension/retraction speed of branches and the portions of extending, retracting, or stationary phase from (K) are quantified. The numbers in the columns indicate quantified ipsilateral exploring branches, contralateral exploring branches, early post-innervation branches, and late post-innervation branches. The extension/retraction speeds were calculated by dividing the total length increase/decrease by the total extension/retraction time during the 14.5 min fast scanning window. Statistics between different groups of branches were done using one-way ANOVA. Statistics between the extending and retracting time fraction within the same group of branches in (J) were done using paired t-test. N.S., P > 0.1; **, P<0.01; ***, P<0.001. (K) Each column indicates the growth status of a single branch across a 14.5-min fast scanning session, from top (beginning) to bottom (end). Each block indicates a 30-sec period. Branches that do not exist for the entire 14.5-min session are only partially represented along the top-bottom axis. Horizontal bars on top of the columns mark branches from the same axon terminals. In post-innervation branches, cyan bars indicate branches from the same axon in the early and late stages, so are orange bars. See Figure S6 for related data.

The enhanced spatiotemporal resolution of AO-LLSM confirmed that ipsilateral innervation occurred via interstitial branching from the contralaterally-projecting axon after the growth cone passed the glomerulus for most ORN types (Figure S6A, green in Figure S6C, D). In the few cases where the ipsilateral branch appeared within the same time period as the contralateral-projecting axon passed through the eventual branching point in our two-photon imaging study (*i* = 0 in Figure 3C), we could determine that the growth cone first innervated the glomerulus, followed by a collateral growing towards the midline that eventually innervated the contralateral antennal lobe (Figure S6B, red in Figure S6C, D). Thus, interstitial branching appears to be a universal rule of ORN axon branching.

When extending along the surface of the antennal lobe, ORN axon terminals usually exhibited simple, needle-like shape. However, we observed a unique multi-branched terminal structure when ORN axons approached their targets (Figure 4D, E; Movie 4). We captured this multi-branched terminal in about 40% ORN axons that were imaged with the appropriate developmental stages in both ipsilateral and contralateral antenna lobes, belonging to multiple ORN types (Figure S6E). Most observations in ipsilateral antennal lobe belonged to ORN types with relatively short waiting time of producing the ipsilateral branch (*i* < 5 in Figure 3C). Typically, 2–5 branches extended from a single axon (Figure S6F), which exhibited dynamic extension or retraction as the branches grew towards the target (see below). We named these “exploring branches”. The exploring branches existed transiently before the axon reached the target (Figure 4D–F, left panels; arrows denote the exploring branches while arrowheads denote the hub from which branches emanate) and disappeared soon after the axon terminal reached the target glomeruli. At the target glomeruli, ORN axon terminals exhibited shorter branches we named “post-innervation branches” (Figure 4F, right panels). Compared to post-innervation branches, exploring branches had longer lifetime (Figure 4G).

The high temporal resolution of AO-LLSM enabled us to analyze the dynamic features of each exploring branch or post-innervation branch. We selected strongly labeled axons and measured the length of each branch across time. Exploring branches had larger maximal length than post-innervation branches (Figure 4H). We categorized each 30-sec imaging interval as extending or retracting if it was part of a time window when the branch continuously extended or retracted to more than 0.5 μm. Otherwise, we defined it as stationary. We separately quantified exploring branches (in both the ipsilateral and contralateral antennal lobes), early post-innervation branches (the first 14.5 minutes imaging period after initial glomerular innervation), and late post-innervation branches (usually 4–6 hours after early post-innervation branches) to assess if terminal dynamics change as the axons mature. The speed of extending and retraction, at 1.5–2 µm/min, appeared similar for both branches in all stages (Figure 4I). However, exploring branches spent more time extending or retracting than post-innervation branches, resulting in fewer stationary periods (Figure 4J, K). Furthermore, extension and retraction in different branches of the same axon terminals appeared to be independent of each other (Figure 4K), consistent with the notion that the behavior of each branch is determined by the microenvironment it samples. The late post-innervation branches exhibited the largest stationary periods, suggesting a trend of being more stable as the axon terminal branches mature during targeting. While the exploring branches exhibited comparable extending and retracting periods, the post-innervation branches exhibited more retracting periods as they matured (Figure 4J). In rare cases when we captured the retraction of most branches within one fast imaging period, we found that different branches were retracted asynchronously (Figure S6G).

In summary, AO-LLSM imaging allowed us to identify a unique exploring branch structure prior to glomeruli innervation. Analysis of branch dynamics across different developmental stages revealed a shift of axon terminal branches from exploration mediated by both branch extension and retraction to a retraction-dominated pruning.

### Cytoskeletal organizations of ORN axon terminals differ from those of cultured neurons

Given the novel structure and properties of the exploring branches, we next examined their cytoskeletal organization and compared with that of the post-innervation branches. According to the textbook depiction, the periphery or leading edge of the growth cone comprises finger-like protrusions called filopodia based on bundled F-actin, and flat, sheet-like protrusions called lamellipodia based on meshwork of F-actin; microtubules fill the axon shaft that usually terminate at the center of the growth cone (Figure 5I, right) (Sanchez-Soriano et al., 2010; Dent et al., 2011). Although microtubules occasionally extend to the periphery of the growth cone, filopodia usually do not contain microtubules (Dent et al., 2011).

**Figure 5.**
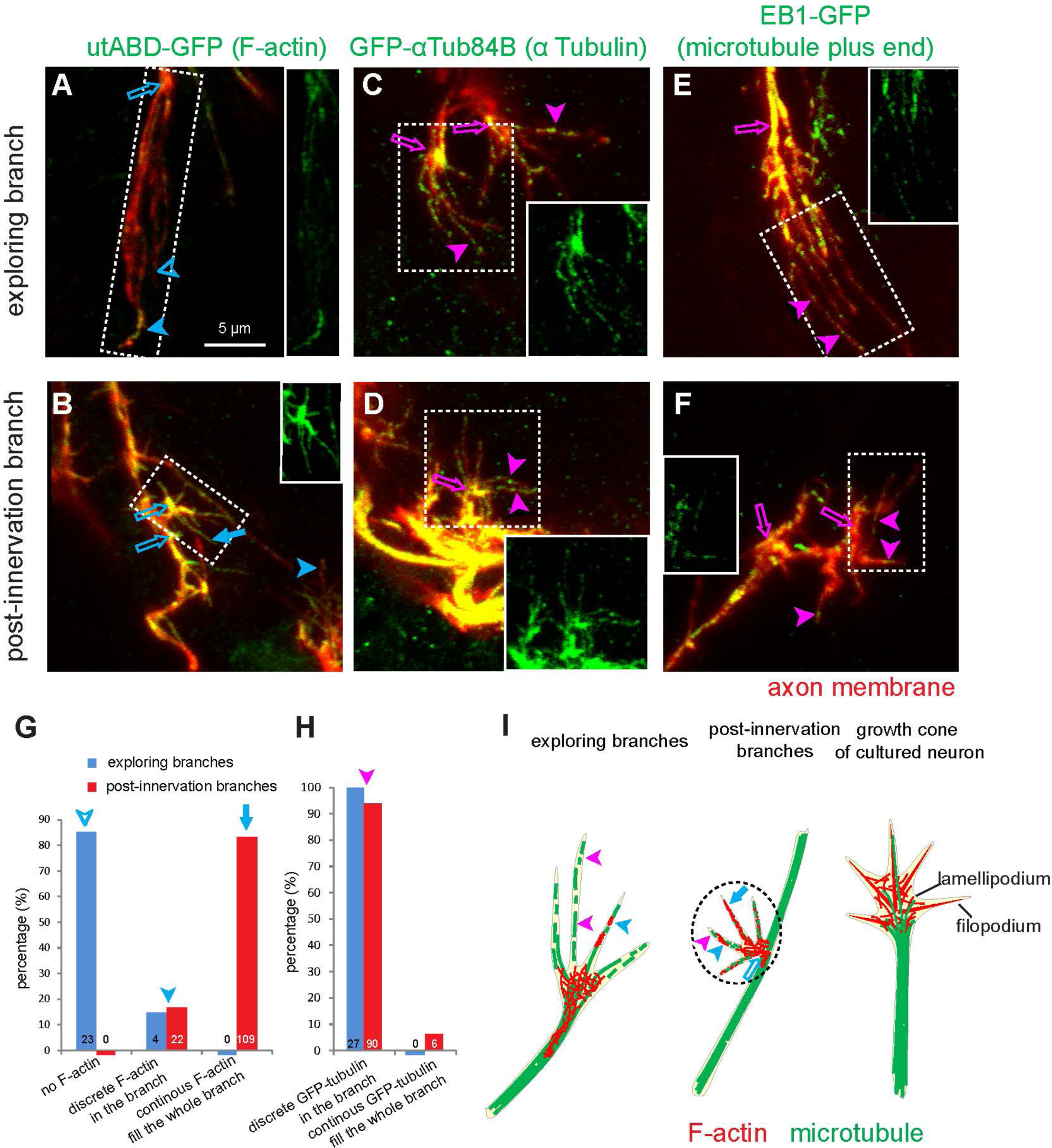
The cytoskeletal organization of ORN axon terminals. (A, B) Max intensity projections of confocal images of ORN axon terminals in fixed pupal brains from 34–36h APF labeled by *pebbled-GAL4*-based MARCM clones co-expressing membrane-Halo (red) and GFP-utABD and visualized with JF-Halo-646 ligand and anti-GFP antibodies, showing the organization of F-actin in exploring branches (A) and post-innervating branch (B). Open arrows denote the hub from which terminal branches extend. Solid arrowheads denote terminal branches with discrete F-actin signal. Blue open arrowheads point branches without F-actin. Solid arrow in (B) denotes a terminal branch filled with F-actin throughout. (C–F) Max intensity projections of confocal images of ORN axon terminals in fixed pupal brains from 34–36h APF labeled by *pebbled-GAL4*-based MARCM clones co-expressing membrane-Halo (red) and GFP-αTub84B (green in C, D) or EB1-GFP (green in E, F) and visualized with JF-Halo-646 ligand and anti-GFP antibodies, showing the organization of microtubules in exploring branches (C, E) and post- innervating branch (D, F). Solid arrowheads denote discrete microtubule fragments in terminal branches. Open arrows indicate the hubs from which terminal branches extend. Insets are GFP-only channel from the regions denoted by the dashed rectangles. (G, H) Quantification of patterns of the F-actin (G) and microtubule (H) from exploring branches and post-innervation branches. Numbers in the column indicate numbers of branches quantified. (I) Summary drawings of cytoskeletal organization in the exploring branches (left) and post-innervation branches (middle) of ORN axonal terminal, compared to and a typical growth cone from cultured neurons (right). Arrows and arrowheads indicate specific structures explained in (A–F). See Figure S6 for related data.

Deviating from the classic depictions of growth cones, which are mostly derived from studies of dissociated neurons in culture, ORN axon terminals—both exploring branches and post-innervating branches—contained multiple thin protrusions (terminal branches) extending from a hub where the axon shaft ends (Figure 4C–E, 5A–F) with no obvious lamellipodia between terminal branches. To examine the cytoskeletal basis of ORN axon terminals, we produced small MARCM clones of ORNs in which the plasma membrane was labeled by a membrane-targeting Halo-tag with Janelia Fluor 646 Halo-tag ligand in the media for explant culture, and F-actin or microtubules was labeled by genetically encoded markers described below, detected by antibody labeling post fixation. To define exploring branches in fixed brains, we selected multi-branched terminals approaching a similar area in the contralateral antennal lobe as the ipsilateral innervated glomeruli. To define post-innervation branches, we selected terminal branches extending from the end of the collateral branch rather than from the contralaterally projecting axon in the ipsilateral antennal lobe.

To study the F-actin distribution in ORN axon terminals, we expressed the F-actin binding domain of utrophin tagged with GFP (GFP-utABD) (Rauzi et al., 2010). In exploring branches, we found that F-actin was concentrated at the hub from which terminal branches emanate (blue open arrows in Figure 5A, I), in contrast to the enrichment in the periphery of the growth cones from cultured neurons. Indeed, most of the exploring branches did not contain detectable F-actin (blue open arrowheads in Figure 5A, I; quantified in Figure 5G); a few branches contained discrete F-actin signal (blue solid arrowheads in Figure 5A, I; quantified in Figure 5G). By contrast, most post-innervation branches were filled with F-actin throughout the entire branch (blue solid arrows in Figure 5B, I, quantified in Figure 5G), whereas the rest of the post-innervation branches contained discrete F-actin patches (blue solid arrowheads in Figure 5B, I, quantified in Figure 5G).

To study microtubule distribution in ORN axons, we expressed GFP-tagged αTub84B (Grieder et al., 2000) or GFP-tagged microtubule plus end–binding protein EB1 (Rusan and Peifer, 2007). The distribution of both markers suggested the presence of microtubule segments in both exploring branches and post-innervation branches of ORNs (magenta solid arrowheads in Figure 5C–F, I), as shown by discrete GFP-αTub84B signal (Figure 5C, D, quantified in Figure 5H) and multiple EB1-GFP puncta along the branches (Figure 5E, F), including the branch tips (Figure S6H).

In summary, the cytoskeletal organization of ORN axon terminals in the antenna–brain explants appeared to differ substantially from the growth cones of dissociated neurons in culture. The microtubule cytoskeleton appears to be a major component in both exploring and post-innervation branches. The post-innervation branches also contain more F-actin, which is most enriched at the hub from which both exploring and post-innervation branches emanate (Figure 5I).

### Contralateral ORN axons are required for correct ORN axon targeting

Bilaterally symmetric axon targeting occurs widely (Yost, 1998; Corballis, 2009; Swanson, 2011). For example, many neurons in the mammalian neocortex and hippocampus target axons to exactly the same regions in the ipsi- and contralateral hemispheres (van Groen and Wyss, 1990; Lodato et al., 2015). The mechanisms that regulate bilaterally symmetric axon targeting are largely unknown (Lodato et al., 2015). Taking advantage of the explant culture system, we asked whether axon targeting of ORNs from one side (hereafter ipsilateral) requires the presence of ORN axons from the other side (hereafter contralateral) by severing the contralateral antennal nerve at a specific developmental time (Figure 6A). Live imaging of sparsely labeled ORN axons showed that axons from the cut side immediately stopped growth and eventually degenerated, while uncut axons from the same explant continued to grow (Figure 6B).

**Figure 6.**
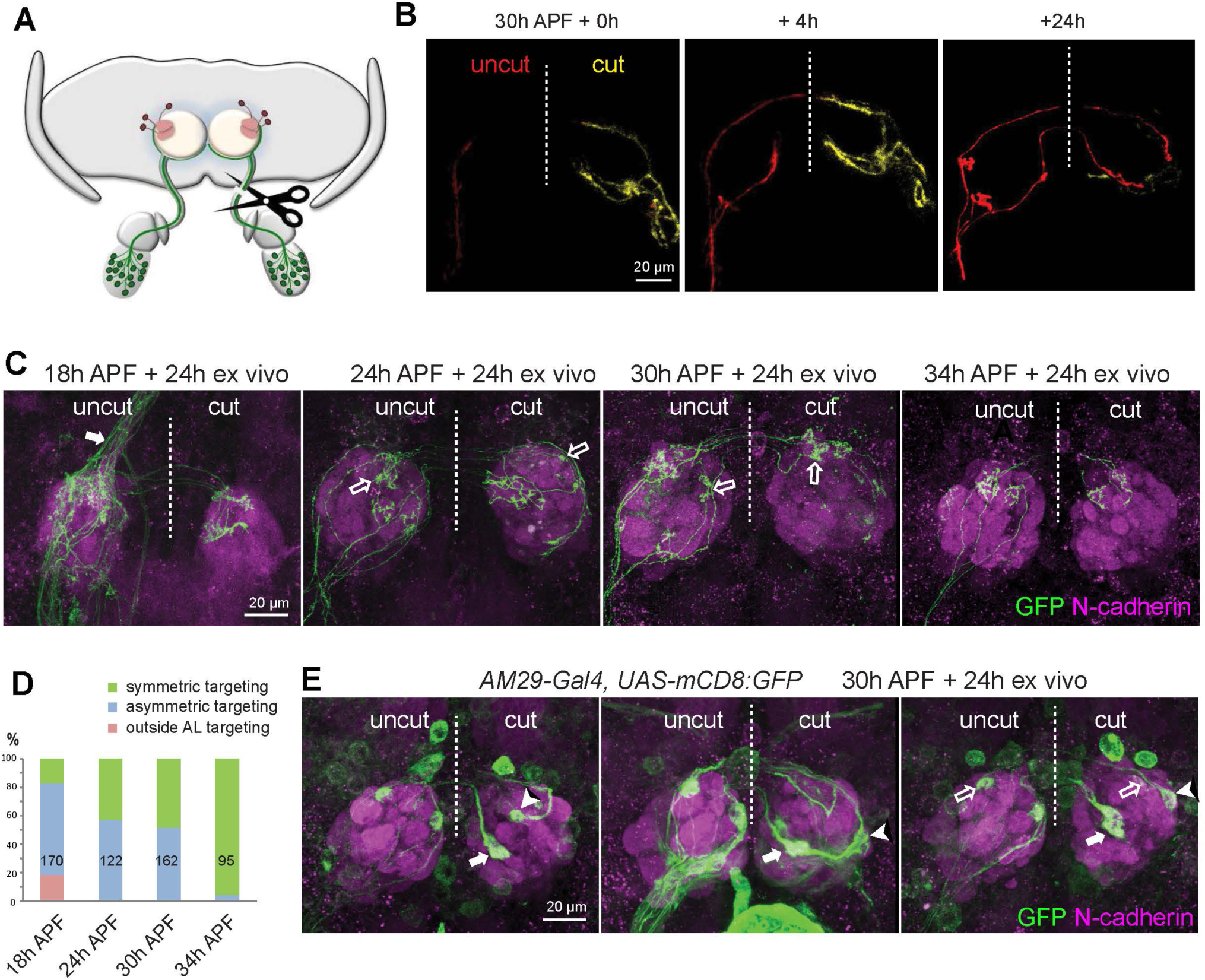
Contralateral ORN axons are required for correct ORN axon targeting. (A) Schematic illustrating unilateral antennal nerve severing prior to explant culture. (B) I*mages* from *time*-*lapse* videos taken at the indicated time points using a two-photon microscope from *pebbled-GAL4/hsFLP; UAS-FRT^100^-stop-FRT^100^-mCD8-GFP/+* explant. Sparsely labeled axons from uncut (red) and cut (yellow) antennae following unilateral antennal nerve severing are shown. Dashed vertical lines indicate midlines, the same as in other images in this figure. (C) Max intensity projection confocal images from four *pebbled-GAL4/hsFLP; UAS-FRT^100^-stop-FRT^100^-mCD8-GFP/+* explants with unilateral antennal nerve severed at 18h, 24h, 30h, and 34h APF, respectively, followed by 24h culture before fixation and staining with anti-GFP and anti-N-cadherin. Solid arrow denotes uncut axons exiting the antennal lobe dorsally. Open arrows denote the asymmetric targeting of single ORN axons in the ipsilateral and contralateral antennal lobes from the same ORN axon. (D) Quantification of fractions of single ORN axons that exhibit targeting outside the antennal lobe (AL), asymmetric targeting, and symmetric targeting of the uncut ORN axons, from explants with one antennal nerve cut at 18h, 24h, 30h and 34h APF (images in C). Numbers in the column indicate the number of axons quantified. (E) Max intensity projection confocal images from *AM29-GAL4,UAS-mCD8-GFP* explants with unilateral antennal nerve severed at 30h APF, followed by 24h culture before fixation and staining with anti-GFP and anti-N-cadherin. Solid arrows and arrowheads mark contralateral mis-targeting of DM6 and DL4 axons respectively. Open arrows mark DM6 and DL4 glomeruli. Mis-targeting of AM29+ ORN axons was observed in 12 out of 14 contralateral antennal lobes. See Figure S7 for related data.

With the sparse labeling of ORNs using the FLPout strategy described above, we initiated explant culture after severing one antennal nerve at 18h, 24h, 30h, and 34h APF. At 18h APF, pioneering ORN axons just reached the antennal lobe (Figure S7A), such that severing of the contralateral antennal nerve would prevent contralateral axons from invading the antennal lobe and thus eliminate bilateral ORN axon–axon interaction. This caused substantial targeting defects of ipsilateral ORN axons, including mistargeting dorsally outside of the antennal lobe (Figure 6C, D). Between 24– 34h APF, ORN axons circumnavigate the antennal lobe, cross the midline, and begin to innervate both antennal lobes (Figure S7A), with increasing opportunities for bilateral ORN axon–axon interactions in the process. We found that contralateral antennal nerve severing at 24h and 30h APF caused ipsilateral ORN axons to target asymmetrically in two antennal lobes (Figure 6C, D), suggesting axon mistargeting on at least one side. However, contralateral antennal nerve severing at 34h APF no longer affected axon targeting specificity of most ipsilateral ORNs on either antennal lobe (Figure 6C, D). Interesting, asymmetric targeting of ipsilateral ORN axons upon severing of the contralateral antennal nerve was observed mostly in ORN types that had a large temporal difference between ipsilateral and contralateral glomerular innervation (Figure S7B, small *i* in Figure 3C).

To determine in which antennal lobe ipsilateral axons mistarget, we examined axon targeting of DM6 and DL4 ORNs in ipsilateral and contralateral antennal lobes upon contralateral antennal nerve severing at 30h APF using driver *AM29-GAL4*. We found that the axons of both ORN types mistargeted only in the contralateral antennal lobe: DM6 ORN axons mistargeted to a stereotyped glomerulus ventrolateral to DM6 glomeruli in the contralateral antennal lobe (solid arrows in Figure 6E), while DL4 ORN axons mistarget to more random glomeruli (solid arrowheads in Figure 6E). Thus, contralateral ORN axons are required for the correct target selection of ipsilateral ORNs in the contralateral antennal lobe.

### ORN axons are required for PN dendrite refinement

Prior to the arrival of ORN axons at the antennal lobe around 18h APF, dendrites of their postsynaptic partner PNs are already present and at least some PN types already target their dendrites to specific regions of the proto-antennal lobe (Jefferis et al., 2004) (Figure 1A). Using the explant system, we first sought to understand how PN dendrites target the correct regions. To label single PNs, we used *GH146- GAL4*-based MARCM and heat shock in a time window when only DL1 single PN clones were labeled (Jefferis et al., 2001) (Figure S8A). Due to the low labeling intensity at early developmental stages, we could only visualize DL1 dendrites starting at ∼23h APF, after they already innervated specific regions of the antennal lobe. We found that at this stage, DL1 dendrites still exhibited active local exploration through terminal branch extension (Figure 7A, arrows; Movie 5). These terminal dynamics gradually decreased after 30h APF when ORNs started to innervate the glomeruli, and the dendritic terminals formed more smooth boundaries (Figure 7B; Movie 5).

**Figure 7.**
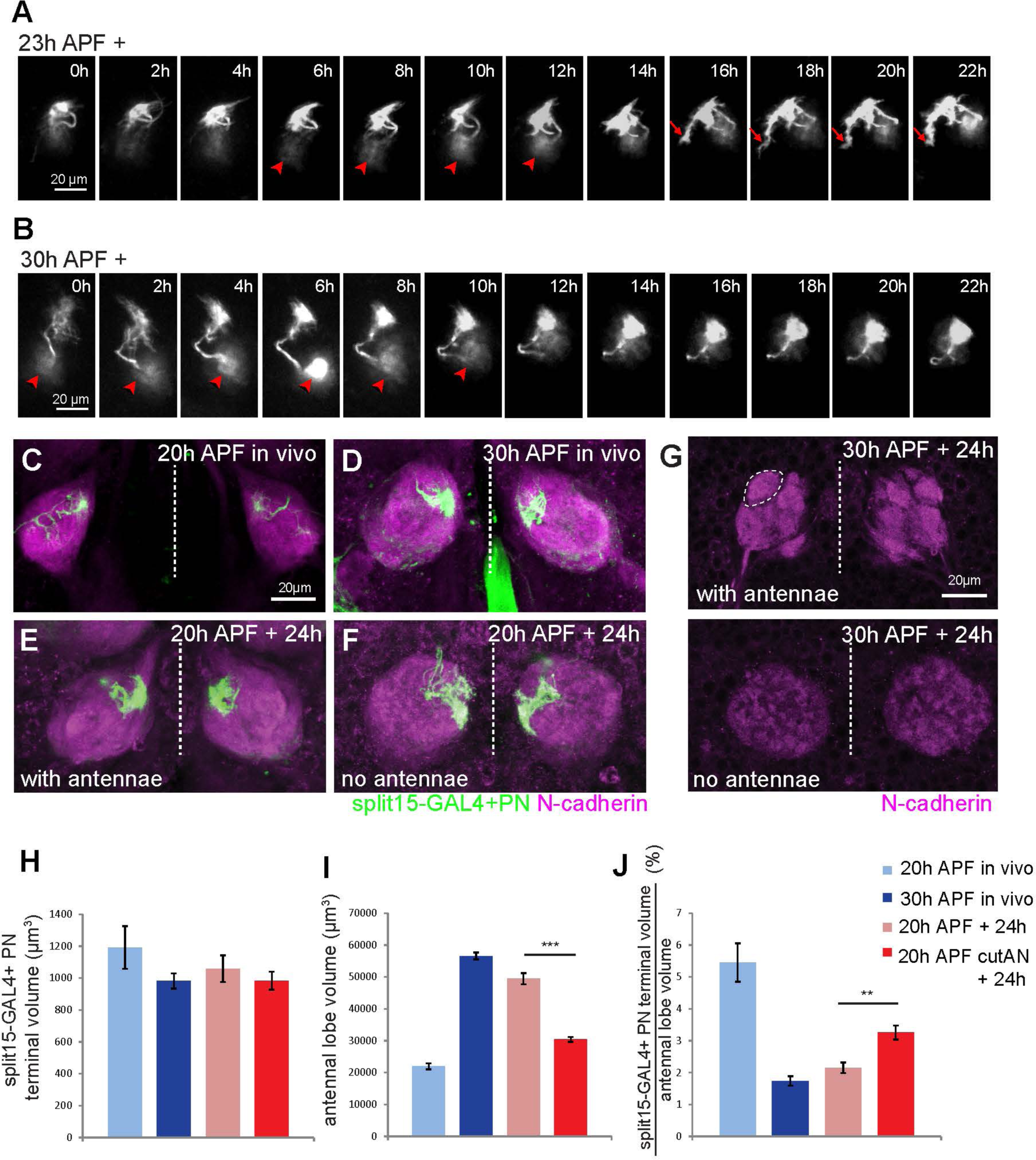
ORN axons are required for PN dendrite refinement. (A, B) *Images* from *time*-*lapse* videos taken from explants dissected at 23h APF (A) or 30h APF (B) using a two-photon microscope, showing dynamics of dendrite terminals across time from a single DL1 PN in each case. Images are partial projection, some of which include the DL1 PN cell body (arrowheads). Arrows denote a dynamic terminal branch exploring local area. Arrowheads mark residual signal from PN cell bodies. (C–F) Max intensity projection confocal images of *split15-GAL4;UAS-mCD8-GFP* brains dissected at 20h APF in vivo (C), 30h APF in vivo (D), and explants dissected from 20h APF in vivo and cultured for 24h with both antennal nerves (AN) intact (E) or cut (F) at the time of dissection. (G) Single confocal image sections of *split15-GAL4;UAS-mCD8GFP* explants cultured for 24h after dissected at 30h APF with antennae nerve intact or severed at the time of dissection. The dashed line marks a newly formed glomerulus. (H–J) Quantification of *split15-GAL4*+ PN dendritic volume (H), total antennal lobe volume (I), and the percentage of *split15-GAL4*+ PN dendritic volume in total antennal lobe (J) from the four conditions indicated, corresponding to panels C–F. Quantifications were done using one-way ANOVA. **: P<0.01, ***: P<0.001. See Figure S8 for related data.

Although PN dendrites innervate the antennal lobe earlier than the ORN axons, there is a window of ∼20 hours between when ORN axons innervate the antennal lobe and when ORN–PN partner matching is complete (Figure 1A). Does PN dendrite development depend on ORN axons? To address this question, we severed both antennal nerves at 20h APF to ablate all ORN axons and compared the targeting of PN dendrites to those with intact antennal nerves (Figure S8A). We assessed targeting of a small subset of PN types labeled by two split GAL4 lines (Yoshi Aso, unpublished data; Xie et al., 2021) (Figure S8B). We found that PN dendrites remained in the same coarse regions of the antennal lobe after 24 hours in explant culture in the absence and presence of the ORN axons (Figure 7E, F; Figure S8C), suggesting that the maintenance of PN dendrites in specific regions of the antennal lobe does not require ORN axons.

However, when ORN axons were ablated, the glomeruli did not form (Figure 7G), consistent with a previous study (Hong et al., 2009). We also observed that PN dendrites occupied larger areas compared to explants with intact ORN axons (Figure 7E, F; Figure S8C). To quantify this effect, we measured the volumes *split-GAL4-15*+ PN dendrites occupied under 4 conditions: 20h APF and 30h APF *in vivo* (Figure 7C, D), 20h APF + 24h explant culture in the presence or absence of ORN axons (Figure 7E, F; quantified in Figure 7H–J). While the volumes of *split-GAL4-15*+ dendrites remained similar at 20h and 30h in vivo (Figure 7H, light and dark blue bars), the antennal lobe volume increased by nearly 3-fold (Figure 7I); as a result, *split-GAL4-15*+ PN dendrites occupied a smaller fraction of the antennal lobe volume (Figure 7J). In other words, PN dendrites were refined as ORN axons invaded the antennal lobe. When the explants were cultured in the presence of ORN axons, a similar expansion of the antennal lobe volume and refinement of PN dendrites occurred after 24h culture (Figure 7H–J, light red bars). However, when the explants were cultured in the absence of ORN axons, antennal lobes no longer expanded, and PN dendrites still occupied the same proportion of the antennal lobe volume after 24h culture (Figure 7H–J, dark red bars). These experiments indicate that ORN axons are required for PN dendrite refinement to a proportionally smaller region of the antennal lobe.

## DISCUSSION

Time-lapse imaging can reveal new insights in complex developmental processes. Here we developed an antennae–brain explant system that mimics olfactory circuit assembly in vivo and is suitable for live imaging study. Systematic imaging of a majority of ORN types using a combination of two-photon microscopy and AO-LLSM—the largest number of identified neuron types in time-lapse imaging studies to our knowledge—in normal and antennal nerve severed samples provided new insights into the cellular basis of the olfactory circuit assembly.

### Cellular mechanisms of target selection

Prior to this study, it is unclear the cellular mechanisms by which ipsilateral target selection is achieved. Among the three possible mechanisms (Figure S5A), our data strongly support the third mechanism: ORN axons send out transient collaterals at multiple locations along the main trunk; the branch that reaches the eventual target becomes stabilized, and further interstitial branches are suppressed (Figure 3E–G). This mechanism accounts for the strong correlation between the position of the ipsilateral branching point and the glomerular position along the VL–DM axis (Figure 3D), the main axis along with ORN axons grow in the ipsilateral antennal lobe. Topographic targeting of retinal ganglion cell axons in the mouse and chick is achieved through preferential arborization of appropriately positioned branches and elimination of ectopic branches (Yates et al., 2001; McLaughlin and O’Leary, 2005), suggesting that the mechanism of transient interstitial branching followed by stabilization applies to the formation of both continuous and discrete neural maps.

The exploring branches we discovered using AO-LLSM imaging (Figure 4) suggest a means by which a growing ORN axon might increase the chance of identifying its target. These exploring branches consist of long, microtubule-based parallel branches that are extending and retracting rapidly and independently, allowing them to sample a relatively large regions for possible targets. The transient occurrence of exploring branches when ORN axons approach their target region suggest that these exploring branches are induced by local cues near target glomeruli to facilitate their target selection. In the ipsilateral antennal lobe, exploring branches were found in ORN types that form ipsilateral branches shortly after the main axon passes by the eventual branching point (small *i* in Figure 3C), consistent with them serving as the precursor to the eventual ipsilateral branch. Exploring branches are also found in axon terminals in the contralateral antennal lobe in ORN types with a wide range of *i* (compare Figure 3C and Figure S6E), suggesting a general role in facilitating contralateral target identification.

For ORN types that have a long delay in extending the ipsilateral branch (large *i* in Figure 3C), we did not observe exploring branches, suggesting a distinct mechanism for consolidating the ipsilateral branch. Nevertheless, we observed dynamic collaterals occurring over a prolong period of time until the formation of the ipsilateral branch, suggesting that these ORN types also use stabilization of transient collaterals as a means to consolidate the ipsilateral branch.

In summary, after the initial trajectory choice such that ORN axons navigate in the half of the antennal lobe where their eventual targets are (Joo et al., 2013) (Figure 1A), we propose that the next critical step in ORN axon development is the stabilization of transient collateral branches by target-derived cues, aided at least in part by the exploring branches. Together, these cellular mechanisms begin to explain how each ORN chooses one of 50 glomerular targets precisely.

### Cytoskeletal organization of axon terminals

A surprising finding is that the cytoskeletal organization of ORN terminals differs substantially from that of classic growth cones, which consist of F-actin-based filopodia and lamellipodia at the periphery and a microtubule-enriched central hub (Figure 5I, right). The terminal branches of ORN axons, in particular the exploring branches which emerge during axon growth near the target, are filled with microtubules, whereas F-actin is concentrated in the central hub (Figure 5). This is unlikely due to species difference, because the classic growth cone cytoskeletal organization is found in neurons (mostly dissociated in culture) from *Aplysia*, *Drosophila*, and mammals (Lin et al., 1994; Sanchez-Soriano et al., 2010; Dent et al., 2011). Indeed, these differences make it difficult for us to define what a growth cone is in ORN axons. We cannot rule out the possibility that F-actin is present in low amount at the terminal of each exploring or post-innervation branch, and it is beyond the detection limit of our utrophin-based F-actin labeling system; if so, each terminal branch would have its own growth cone at its tip that resembles classic growth cones. Even if that were the case, ORN axon terminals still differ from classic growth cones by having multiple microtubule-based parallel branches emanating from an F-actin rich central hub. Indeed, EB1-GFP puncta can be found at the tip of the branch (Figure S6H), suggesting that microtubules can fill the entire branch. Microtubule polymerization has been shown to mediate membrane extension directly in lipid vesicle (Fygenson et al., 1997).

We suspect that the deviation of cytoskeletal organization in ORN axon terminals from the classic growth cone is likely due to the much more complex environments axon terminals need to explore in the brain compared with the primary culture. Our findings have important implications for signaling mechanisms that convert cell-surface recognition into cytoskeletal-based structural changes in axon terminals during axon targeting. Specifically, we suggest that regulation of the microtubule cytoskeleton might be particularly important at initial stages of target selection.

### Bilateral interactions in bilaterally symmetrical connections

Bilaterally symmetric organization of the nervous system is a cardinal feature of all bilaterians (Swanson, 2011). Despite the prevalence of bilaterally symmetric axon targeting, little is known about the underlying mechanisms. Our unilateral antennal nerve severing provides, to our knowledge, the first evidence for bilateral interactions in target selection. The simplest cellular mechanism is direct interactions between ipsilateral and contralateral ORN axons. These interactions may facilitate midline crossing, disruption of which may cause some axons to leave the antennal lobe instead (Figure 6C). At a later developmental stage, bilateral axon-axon interactions between the same ORN type may facilitate target selection of contralateral ORNs. Our data does not rule out the possibility that bilateral interactions may be indirect; for example, ipsilateral ORNs may change the properties of their partners PNs, which in turn regulate target selection of contralateral ORNs. Interestingly, upon unilateral antennal nerve severing, the asymmetric targeting from the uncut ORNs was mostly found in the ORN types that sequentially innervate ipsilateral and contralateral glomeruli (Figure S7B). The early ipsilateral innervation of these ORNs may be important for the contralateral targeting of the same ORN type from the other side. The ease of disrupting bilateral interactions in explant cultures provides a means to further investigate cellular and molecular mechanisms of bilateral interactions.

In conclusion, time-lapse imaging has greatly enriched our understanding of the cellular events that enable the step-wise assembly of the fly olfactory circuit, and highlight the precise genetic control of multiple steps during ORN axon targeting. These include the choice of a trajectory along which an ORN axon navigates the ipsilateral antennal lobe, the timing and location of stabilizing its ipsilateral branch, as well as the interactions with contralateral ORN axons to cross the midline and innervate its contralateral target. Finally, ORN axons also help refine dendrites of their partner PNs, which pattern the antennal lobe first. The stage is set to combine live imaging and the cellular insights it has brought with genetic manipulations of key wiring molecules identified by genetic, transcriptomic, and proteomic approaches to reach a new level of mechanistic understanding of the circuit assembly process.

## AUTHOR CONTRIBUTIONS

T.L. and L.L. designed experiments. T.F. and T.L. did the AO-LLSM imaging experiments under the guidance of E.B. H.L. screened PN split GAL4 lines and Q.X. assisted fly experiments. D.J.L. helped generating transgenic flies. T.L. performed all the rest experiments and analyzed the data with discussion with L.L. T.L. and L.L. wrote the paper, with input from all other coauthors. L.L. supervised the project.

## ACKNOWLEDGMENTS

We thank M. Ozel and R. Hiesinger for their advice on the explant culture; T. Lecuit, L. Lavis, and Bloomington *Drosophila* Stock Center for reagents; M. Wagner for technical help of the two-photon microscopy; Y. Ge for assistance on fly work; J. Li, C. McLaughlin, D. Friedmann, Y. Takeo, M. Wagner, K.K.L. Wong, J. Baruni, D. Pederick, D. Milkie, G. Upadhyayula, L. Lavis, T. Brown, P. Dong, K. Shen, Z. Li, R. Zhong for discussions; T. Lee for sharing experimental equipment at Janelia Research Campus; H.L. was a Stanford Neuroscience Institute interdisciplinary postdoctoral scholar. E.B. and L.L. are Howard Hughes Medical Institute (HHMI) investigators. This work was supported by National Institutes of Health grants 1K99DC01883001 (to T.L.) and R01-DC005982 (to L.L.).

## STAR METHODS

### KEY RESOURCES TABLE

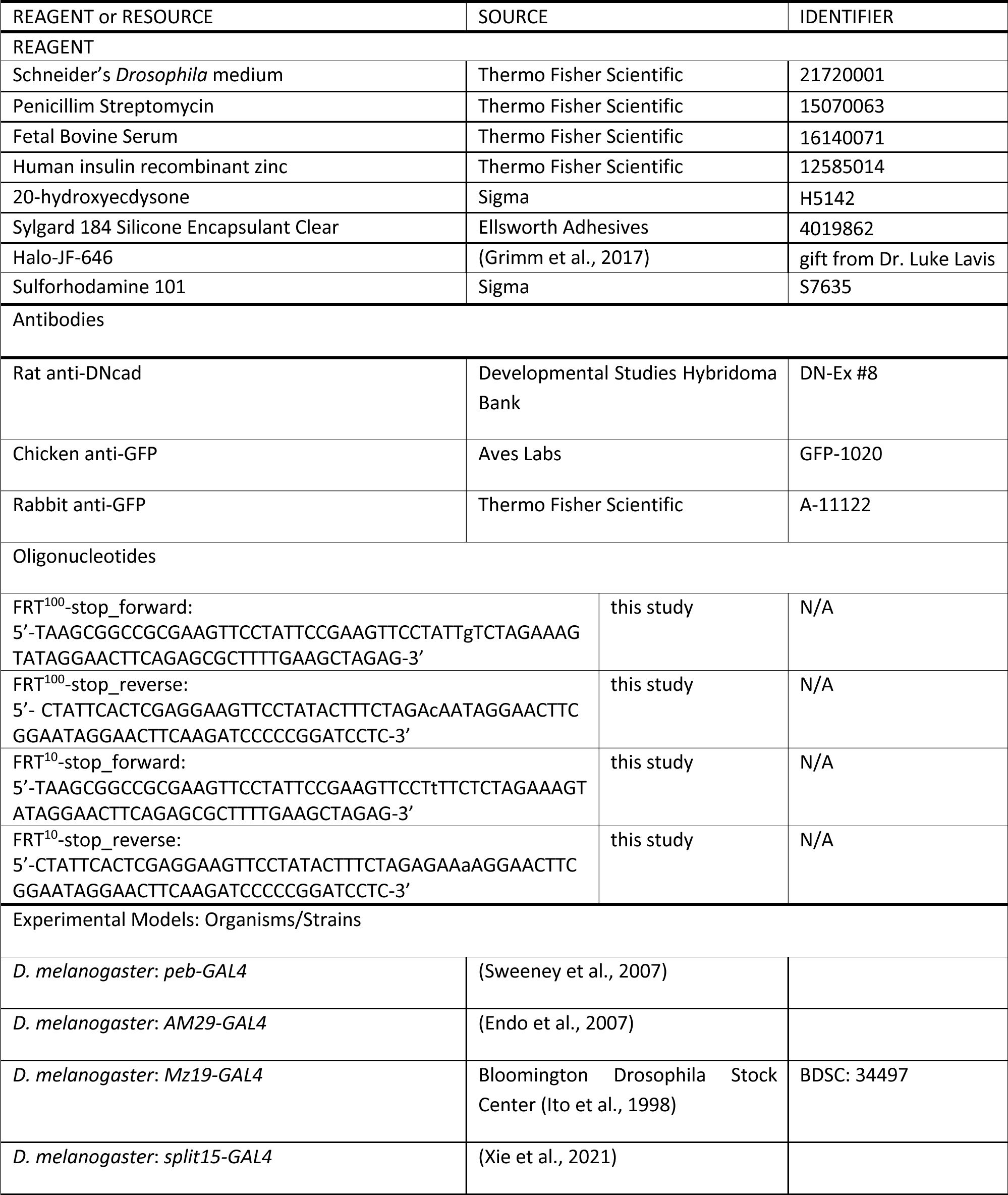
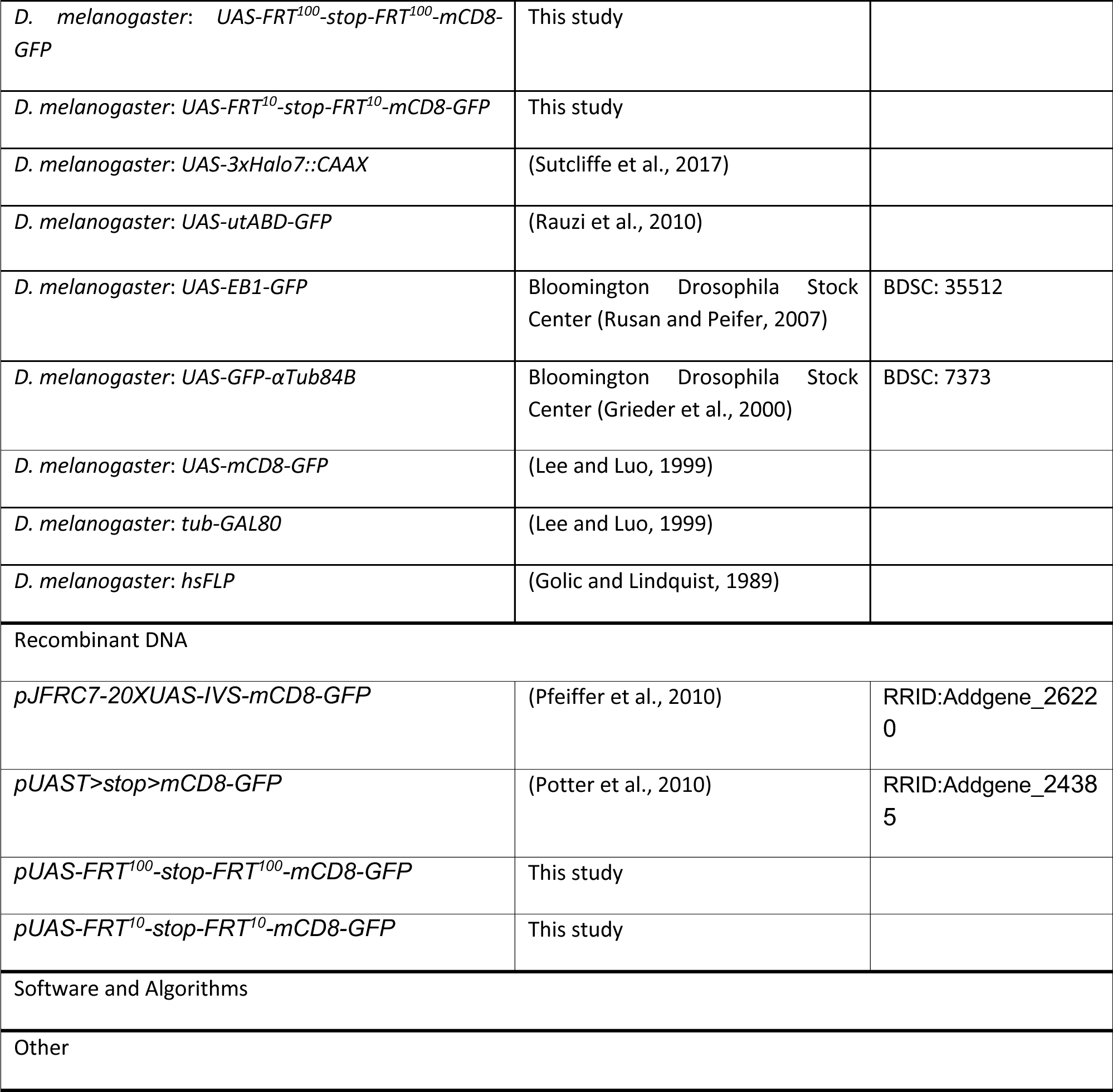

Further information and requests for resources and reagents should be directed to the Lead Contact, Liqun Luo. All unique reagents generated in this study are available from the Lead Contact.

### EXPERIMENTAL MODEL AND SUBJECT DETAILS

#### *Drosophila* stocks and genotypes

Flies were raised on standard cornmeal medium with a 12h/12h light cycle at 25°C. Complete genotypes of flies in each experiment are described in Table S1.The following lines were used:*pebbled-GAL4* (pan-ORN GAL4)(Sweeney et al., 2007), *AM29-GAL4* (DM6 and DL4 ORN GAL4)(Endo et al., 2007), *Mz19-GAL4* (DA1, VA1d, and DC3 PN GAL4) (Ito et al., 1998), *split15-GAL4* (Yoshi Aso, unpublished data; Xie et al., 2021), *UAS-FRT^100^-stop-FRT^100^-mCD8-GFP*, *UAS-FRT^10^-stop-FRT^10^-mCD8-GFP* (this study), *UAS-3xHalo7::CAAX* (Sutcliffe et al., 2017), *UAS-utABD-GFP* (Rauzi et al., 2010), *UAS-EB1-GFP* (Rusan and Peifer, 2007), *UAS-GFP-αTub84B* (Grieder et al., 2000), *UAS-mCD8-GFP*, *tubP-GAL80* (Lee and Luo, 1999), *hsFLP* (Golic and Lindquist, 1989).

### METHOD DETAILS

#### Explant dissection and culture

The 0h APF pupae were identified by the white color of the cuticle of pupae. These pupae were heat shocked in 37°C water bath for different amount of time to induce sparse clones (details indicated in elsewhere) and aged at 25°C for certain hours before dissection. Dissection was done in pre-cooled Schneider’s insect media (Thermo Fisher Scientific) with Penicillim Streptomycin (Thermo Fisher Scientific). The external brown shells of the pupae were first removed using forceps. The micro dissection scissor was used to cut the trunk and remove the semi-transparent cuticle covering the dorsal and ventral sides of the brain. A small piece of cuticle attached to two retinae was left to hold the two antennae with the brain. Fat body particles were then cleaned through gentle pipetting. At this point the antennal nerves that connect the two antennae to the brain are visible and can be severed by the micro scissor in certain experiments (Figures 6, 7). After severing the antennal nerve(s), the antenna(e) were left attached to the small piece of cuticle during culture (Figure 6, 7). Following dissection, the antennae-brain explant was transferred to a culture dish with silgard layer on the bottom and 500 μl culture media. Two micro pins were used to pin the explant to the silgard through the two optic lobes. The silgard plate with explant was then carefully moved to the imaging station. 10 ml culture media was added gently to the plate before imaging. The culture condition for explant was modified from Ozel et al., 2015, containing: Schneider’s insect media (Thermo Fisher Scientific), 10% Fetal Bovine Serum (Thermo Fisher Scientific), 10 μg/ml human insulin recombinant zinc (from 4mg/ml stock solution, Thermo Fisher Scientific), 1 μg/ml 20-hydroxyecdysone (from 1 mg/ml stock solution in ethanol, Sigma), and Penicillin/streptomycin (1:100 from stock: 10000 IU/ml penicillin, 10 mg/ml streptomycin, Thermo Fisher Scientific). The culture media was oxygenated by pumping oxygen for ∼ 30 min before use.

#### Generation of sparse labeling FLPout reporter flies

To generate FLPout reporters *pUAS-FRT^100^-stop-FRT^100^-mCD8-GFP* and *pUAS-FRT^10^-stop-FRT^10^-mCD8- GFP* constructs (Figure 2), we PCR amplified the transcriptional terminator, stop, sequence from *pUAST>stop>mCD8-GFP* (addgene #24385, Potter et al., 2010) and subcloned into *pJFRC7-20XUAS-IVS- mCD8-GFP* (addgene # 26220, Pfeiffer et al., 2010) through NotI and XhoI. *FRT^100^* or *FRT^10^* sequences (mutant *FRT*s with ∼100-fold or ∼10-fold less efficiency compared to wild type *FRT*; Golic and Lindquist, 1989) flanking stop sequence were added through the primers for PCR. Both constructs were sequence verified. The sequences of primers used for PCR reaction are below:

FRT^100^-stop_forward:5’-TAAGCGGCCGCGAAGTTCCTATTCCGAAGTTCCTATTgTCTAGAAAGTATAGGAACTTCAGAGCGCTTTTGAAGCTAGAG

FRT^100^-stop_reverse:5’-CTATTCACTCGAGGAAGTTCCTATACTTTCTAGAcAATAGGAACTTCGGAATAGGAACTTCAAGATCCCCCGGATCCTC

FRT^10^-stop_forward:5’-TAAGCGGCCGCGAAGTTCCTATTCCGAAGTTCCTtTTCTCTAGAAAGTATAGGAACTTCAGAGCGCTTTTGAAGCTAGAG

FRT^10^-stop_reverse:5’-CTATTCACTCGAGGAAGTTCCTATACTTTCTAGAGAAaAGGAACTTCGGAATAGGAACTTCAAGATCCCCCGGATCCTC

Both *pUAS-FRT^100^-stop-FRT^100^-mCD8-GFP* and *pUAS-FRT10-stop-FRT10-mCD8-GFP* transgenes were integrated into *86Fb* landing site (Bischof et al., 2007).

#### Immunocytochemistry

Dissection and immunostaining of fly brains and pupal antennae were performed according to previously described methods (Wu and Luo, 2006; Li et al., 2016). The brains were dissected in PBS (phosphate buffered saline; Thermo Fisher), and then fixed in 4% paraformaldehyde (Electron Microscopy Sciences) in PBS with 0.015% Triton X-100 (Sigma-Aldrich) for 20 minutes on a nutator at room temperature. Explant was fixed in the same condition after culture. Fixed brains were washed with PBST (0.3% Triton X-100 in PBS) four times, each time nutating for 20 minutes. The brains were then blocked in 5% normal donkey serum (Jackson ImmunoResearch) in PBST for 1 hour at room temperature or overnight at 4°C on a nutator. Primary antibodies were diluted in the blocking solution and incubated with brains for 36–48 hours on a nutator at 4°C. After washed with PBST four times, each time nutating for 20 minutes, brains were incubated with secondary antibodies diluted in the blocking solution and nutated in the dark for 36–48 hours at 4°C. Brains were then washed again with PBST four times, each time nutating for 20 minutes. Immunostained brains were mounted with Slow Fade anti-fade reagent (Thermo Fisher) and stored at 4°C before imaging. Primary antibodies used in immunostaining include: rat anti-DNcad (1:40; DN-Ex#8, Developmental Studies Hybridoma Bank), chicken anti-GFP (1:1000; GFP-1020, Aves Labs), rabbit anti-GFP (1:1000, A-11122, Thermo Fisher Scientific). Donkey secondary antibodies conjugated to Alexa Fluor 488/647(Jackson ImmunoResearch or Thermo Fisher Scientific) were used at 1:1000.

#### Image acquisition and processing

Images of fixed brains were acquired by a Zeiss LSM 780 laser-scanning confocal microscope (Carl Zeiss), with a 40x/1.4 Plan-Apochromat oil objective (Carl Zeiss). Confocal z stacks were obtained at 1–2 µm intervals at the resolution of 512x512. Images were exported as maximum projections or single confocal sections by ZEN (Carl Zeiss) in the format of TIFF. Photoshop (Adobe) was used for image rotation and cropping. Two-photon microscopy-based imaging was performed at room temperature using a custom-built two-photon microscope (Prairie Technologies), a Chameleon Ti:Sapphire laser (Coherent) and a 20×water-immersion objective (1.0 NA; Zeiss). For all two-photon microscopy-based imaging, the excitation wavelength was at 920 nm. Z stacks were obtained at 2 µm intervals. The pixel dwell time was 10 µs. Images were exported as maximum projections or single confocal sections by FIJI in the format of TIFF.

For AO-LLSM based imaging, the excitation and detection objectives along with the 25-mm coverslip were immersed in ∼40 ml of culture medium (see in Explant dissection and culture) at room temperature (22 ± 1°C). Explant brains with membrane targeting Halo-JF-646 expressed in sparse ORN clones were immobilized on a thin Sylgard layer (∼2 mm) attached to the surface of the coverslip using two pins and were excited using 642 nm lasers operating at ∼2–10 mW (corresponding to ∼10–50 μW at the back aperture of the excitation objective) with an exposure time of 20–50 msec. Dithering lattice light-sheet patterns with an inner/outer numerical aperture of 0.38/0.4 was used. The optical sections were collected by an axial step size of 250 nm in detection objective coordinate, with a total of 81–201 steps (corresponding to a total axial scan range of 20–50 μm). Emission light from JF-646 was captured by a Hamamatsu ORCA-Flash 4.0 sCMOS cameras (Hamamatsu Photonics, Hamamatsu City, Japan). Prior to the acquisition of the time series data, the imaged volume was corrected for optical aberrations using two-photon “guide star” based adaptive optics method. Each imaged volume was deconvolved in C++ using Richardson-Lucy algorithm on HHMI Janelia Research Campus’ computing cluster with experimentally measured point spread functions obtained from 200 nm fluorescent beads (Thermo Fisher). The AO-LLSM was operated using a custom LabVIEW software (National Instruments, Woburn, MA). Image analysis was performed using FIJI.

#### Sparse genetic labeling and dye labeling

To sparsely label a few ORNs from any types using the FLPout reporters we generated in this study for time-lapse imaging (Figure 2), we collected *hsFLP/pebbled-GAL4;;UAS-FRT^100^-stop-FRT^100^-mCD8-GFP/+* pupae at 0h APF and heat shocked at 37°C for 40 min at 0h APF. These pupae were then aged at 25°C for 30h before explant dissection and two-photon imaging. To sparsely label a few ORNs with membrane targeting Halo-tag using MARCM (Figure 4), we heat shocked *pebbled-GAL4, FRT19A/tub-GAL80, hsFLP, FRT19A;;UAS-3xHalo7::CAAX/+* at 37°C for 30 min 2 days before puparium formation. We then collected these pupae at 0h APF and aged them at 25°C for 28h, followed by explant dissection, incubation with 2 μM Halo-JF-646 (Grimm et al., 2017) in 1 ml oxygenated culture media for 1h at room temperature. These explants were then incubated with 1µM Sulforhodamine 101 (Sigma) in 1ml culture media for 5 min at room temperature followed by two time washing in culture media before AO-LLSM based imaging in 50 ml culture media at room temperature. To induce DL1 PN clones, we heat shocked *yw, UAS-mCD8-GFP, hsFLP; FRTG13, GH146-FLP, UAS-mCD8-GFP/FRTG13, tubP-GAL80* at 37 °C for 1 hour at approximately 0-4 hours after larval hatching (Jefferis et al., 2004).

#### Imaging analysis

To quantify growth of ORN axons across time (Figure 3A, B), we measures the change of ORN axon curve length in the antennal lobe in each 20 min imaging interval and plotted to the timing of midline crossing. In the ipsilateral antennal lobe, we defined the last imaging scan when axons have not entered the midline area as time 0. In the contralateral antennal lobe, we defined the first imaging scan when the axons exit the midline area as time 0. To quantify the ipsilateral branching point (Figure S5A-C), *La* was measured as the curve length from the antennal lobe entry point to ipsilateral branch point or the center of branching area in the main axon. *Lb* was measured from the midline-axon crossing point to the ipsilateral branch point or the center of branching area in the main axon. The curve length of axons was measured using IMARIS. To describe the pattern of ipsilateral branches (Figure 3F, G), we drew the outlines of the branch terminals from the projection images. To analyze the fast dynamics of exploring branches and post-innervation branches, we measured the curve length of each branch across time using imageJ and categorized each 30-sec imaging interval as extending or retracting if it was part of a time window when the branch continuously extended or retracted to more than 0.5 μm. Otherwise, we defined it as stationary. To calculate extending or retracting speed of each branch, we summed up total length increase or decrease during the 14.5 min fast scanning period and divided by the extending or retracting time period, respectively. Statistical analyses were performed using t-test, one-way ANOVA, two-way ANOVA or paired t-test as indicated in the figure legend for each experiment using Excel. The significance in paired t-test was based on P(T≤t) two tails.

---

**Figure S1.**
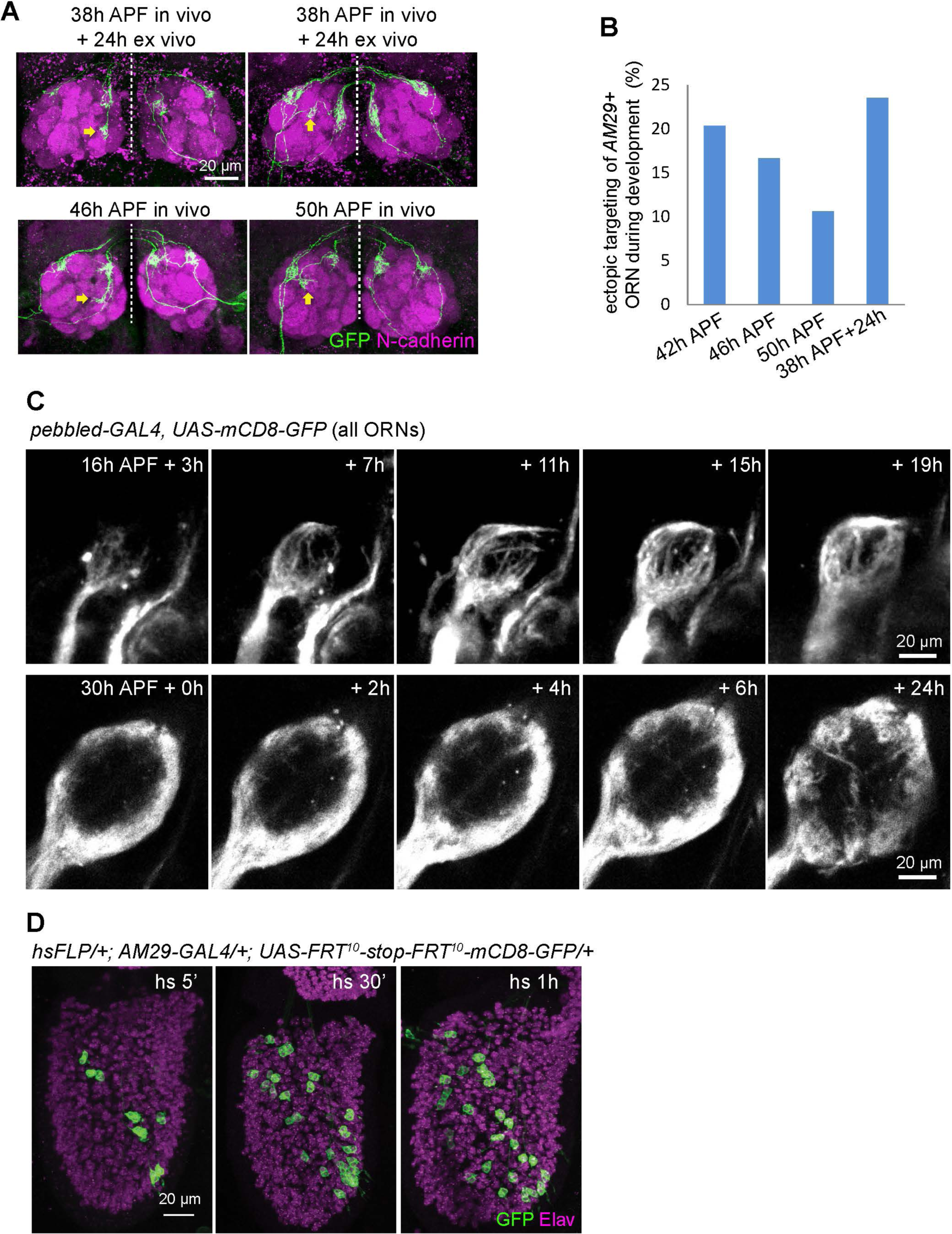
The explant closely mimics normal olfactory circuit development, related to Figure 1. (A) Max intensity projection confocal images from explants (top row) or brains in vivo (bottom row). *AM29-GAL4*+ MARCM clones are labeled by expression of mCD8-GFP. Arrows mark ectopic targeting of *AM29-GAL4*+ ORN axons to ventral or lateral side of DM6 in both explants and in vivo. (B) Quantifications of ectopic targeting of MARCM clones of *AM29-GAL4*+ ORN axons during development in vivo and ex vivo. (C) *Images* from *time-lapse* videos taken by a two-photon microscope from *pebbled-GAL4;UAS-mCD8- GFP* explant at indicated time points. (D) Max intensity projection confocal images from 3^rd^ antennal segment at 48h APF. Different heat shock time induces different number of *AM29-GAL4*+ ORNs by *UAS-FRT^10^-stop-FRT^10^-mCD8-GFP* reporter.

**Figure S2.**
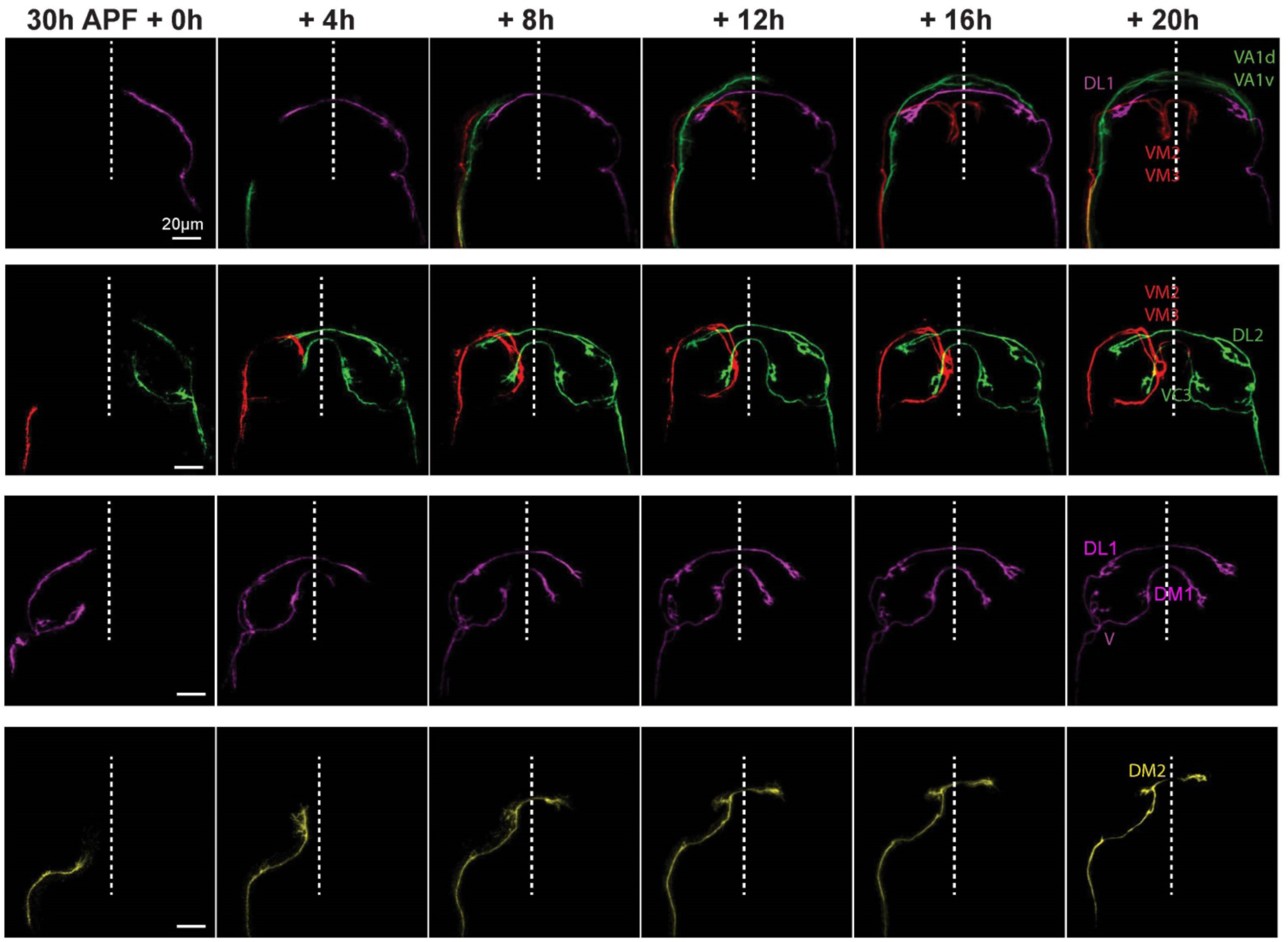
Examples of time-lapse series of single ORN axons during target selection, related to Figure 2. *Images* from *time-lapse* videos from a two-photon microscope at the indicated time points of sparsely labeled ORNs from four samples (one per row). The identity of each ORN was revealed by fixing the explant at the end of the culture and staining with neuropil marker N-cadherin. Different single ORN axons are pseudocolored.

**Figure S3.**
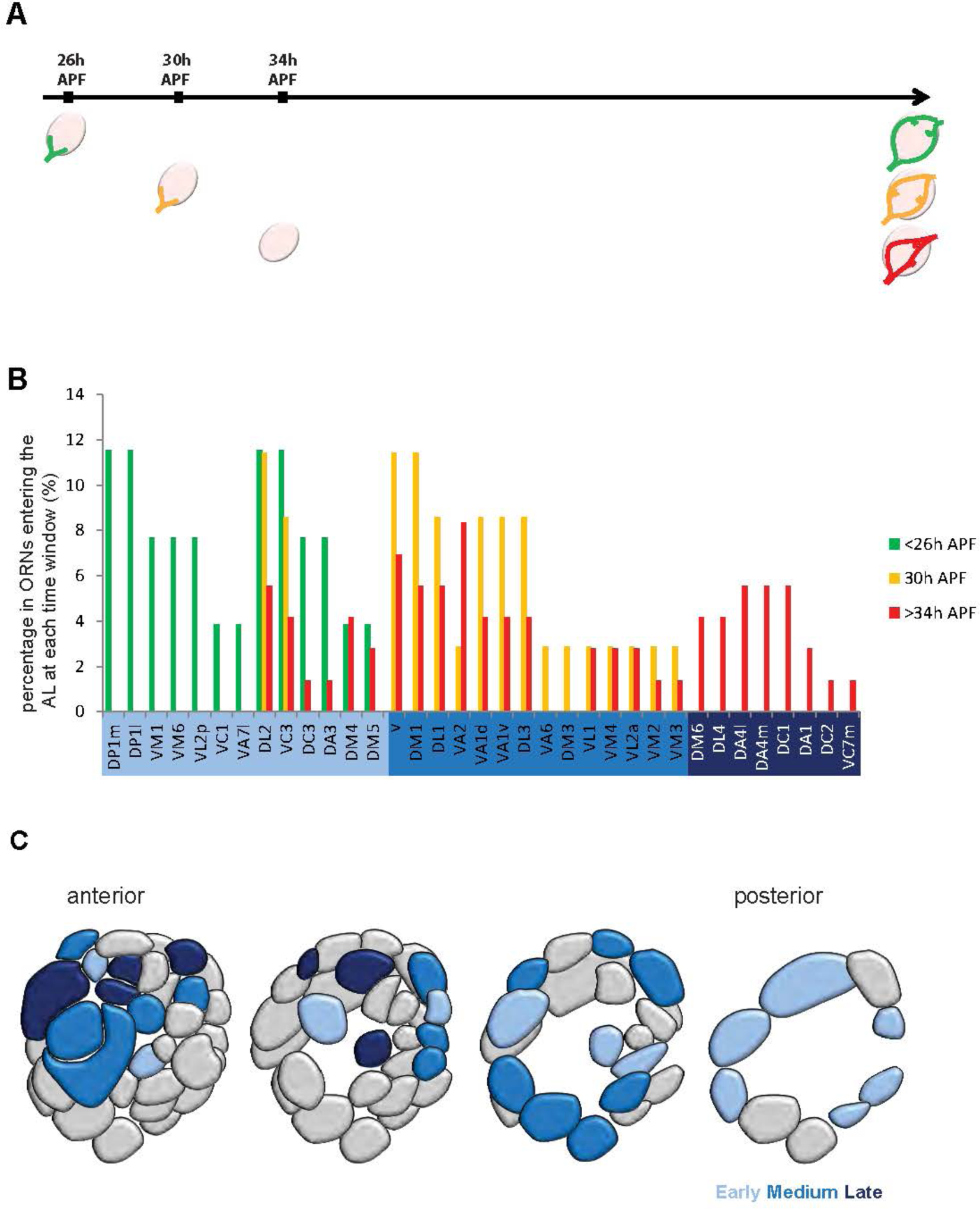
Axons from different ORN types reach the antennal lobe within different time windows, related to Figure 2. (A) ORN axons that enter the antennal lobe at different time windows were identified by dissecting the pupal brains at 26h APF, 30h APF and 34h APF and selected based on the arrival at the antennal lobes, followed immediately by time-lapse imaging. Each ORN axon was tracked by time-lapse imaging and the targeting glomerular identity was determined by fixing and staining at the end of the 48h culture for 26h APF-dissected explants and 24h culture for 30h APF- and 34h APF-dissected explants. (B) Percentage of ORN types in the whole ORN population that enter the antennal lobe at indicated time windows. The ORN types are ordered according to the time of the earliest entering cases. (C) Glomerular map showing the distribution of ORN types with early- (light), middle- (medium) and late- (dark) arriving axons.

**Figure S4.**
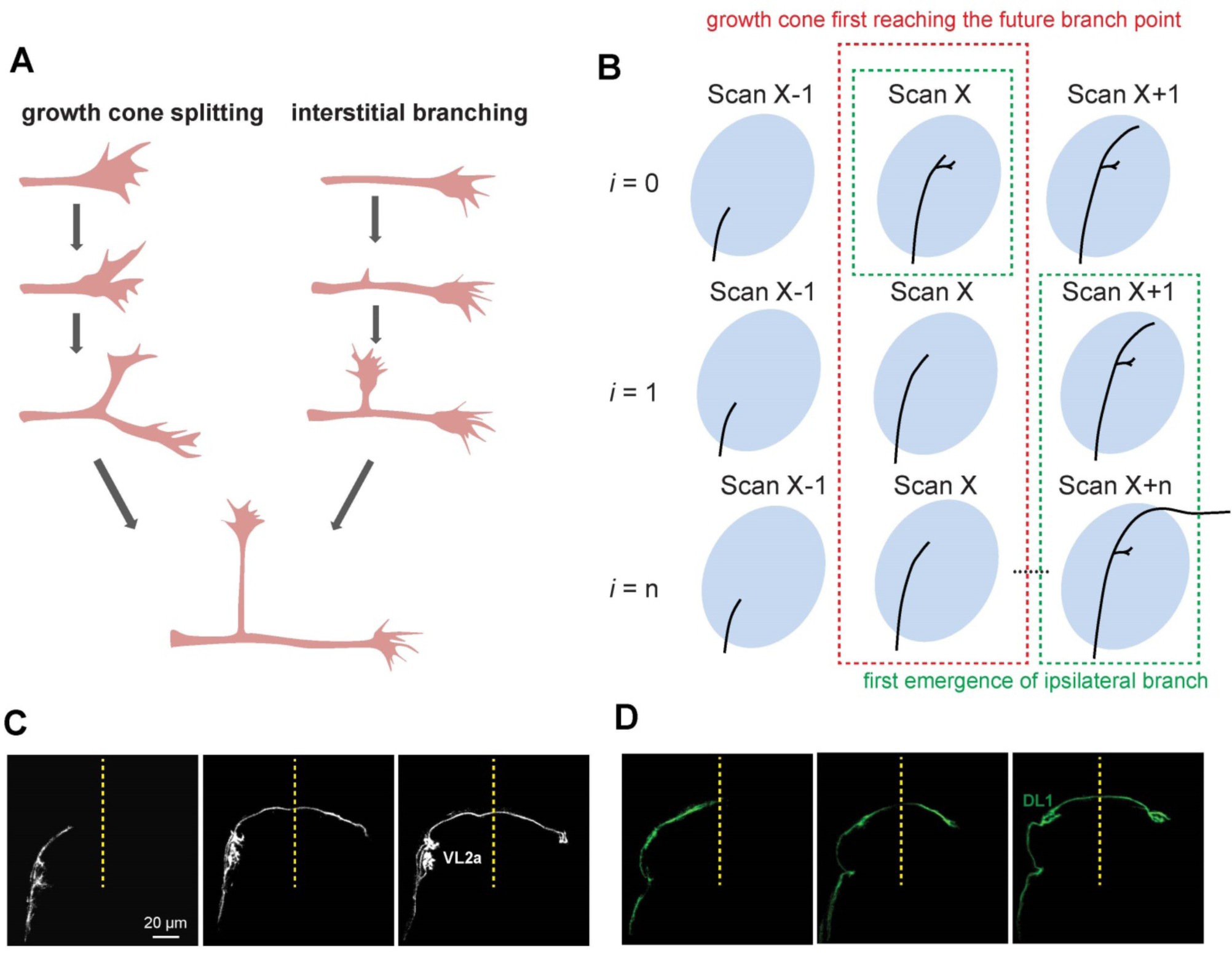
Temporal analysis of ipsilateral innervation of ORN axons, related to Figure 3. (A) Schematic showing two mechanisms of branching: growth cone splitting versus interstitial branching. (B) Illustration of examples when imaging sessions *i* = 0, 1, n in determining axon branching mechanism (Figure 3C). (C, D) Two-photon microscopy-based time-lapse images of VL2a and DL1 ORNs showing examples of ORN types with large timing difference (C, small *i*) and small timing difference (D, large *i*) between ipsilateral and contralateral innervations.

**Figure S5.**
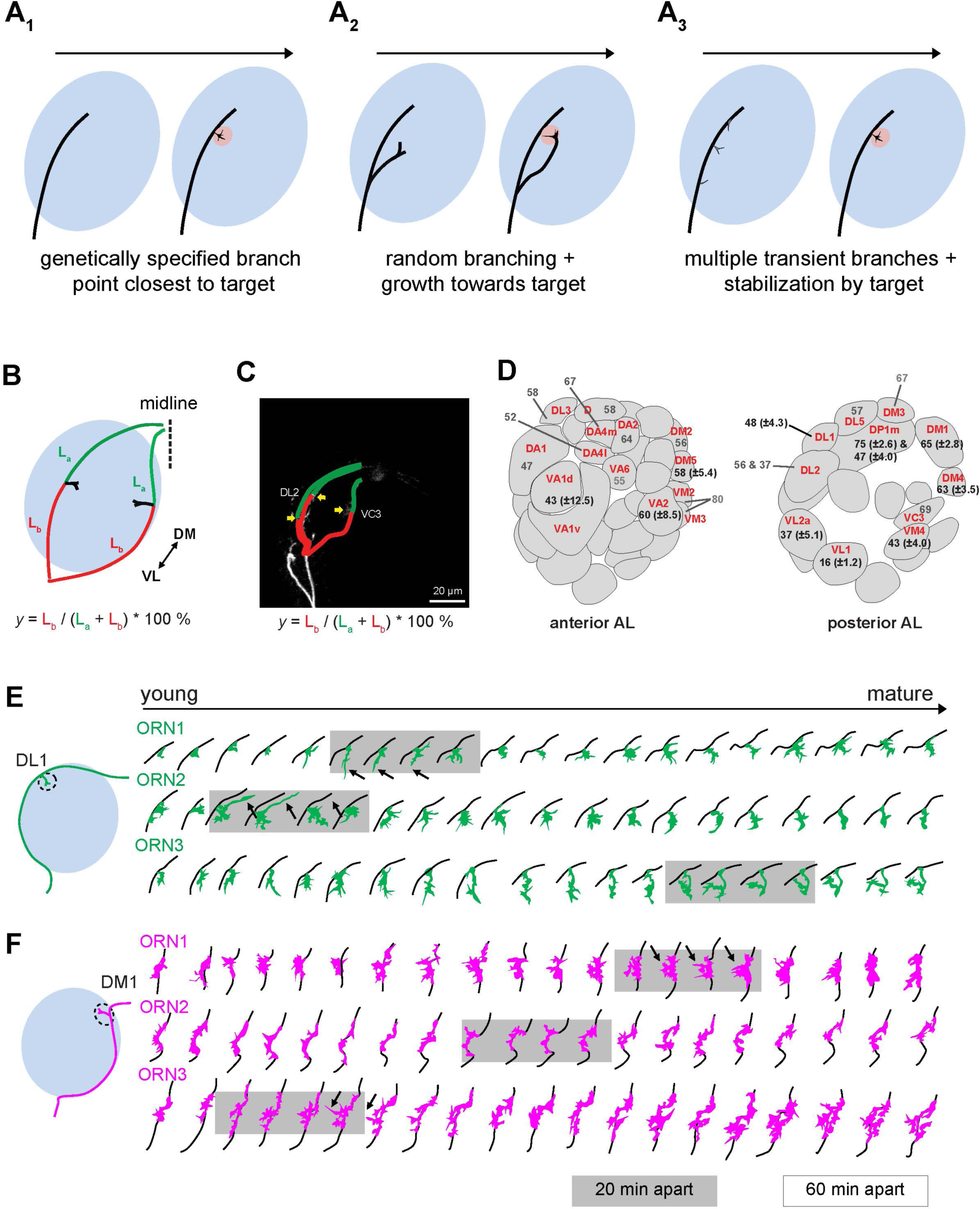
Spatial analysis of ipsilateral innervation of ORN axons, related to Figure 3. (A) Schematic illustrating three mechanisms of ipsilateral innervation: (A1) genetically specified branching point closest to the target; (A2) random branching followed by growth of ipsilateral branch towards the target; (A3) multiple transient branches and stabilization of the branch close to the target . (B) Schematic illustrating the calculation of ipsilateral branch point index *y*. (C) An example of measuring *y* for DL2 and VC3. Two arrows on DL2 mark two ipsilateral branching points. (D) *y* from 90 single ORNs time-lapse-imaged are quantified and shown with their glomerular identities. Numbers in brackets are SEMs from the same ORN type with ≥3 samples. (E, F) Silhouettes showing the shapes of ipsilateral branches from 3 single DL1 (E) and 3 single DM1 (F) ORNs across time. Each row is a single ORN. The main axon shafts are shown in black and ipsilateral branches are shown in colors. Neighboring images are 1 hour apart except those in gray shade, which are 20 min apart. Arrows denote examples of branch retraction. Schematics on the left show the positions of the DL1 and DM1 glomeruli in the left antennal lobe.

**Figure S6.**
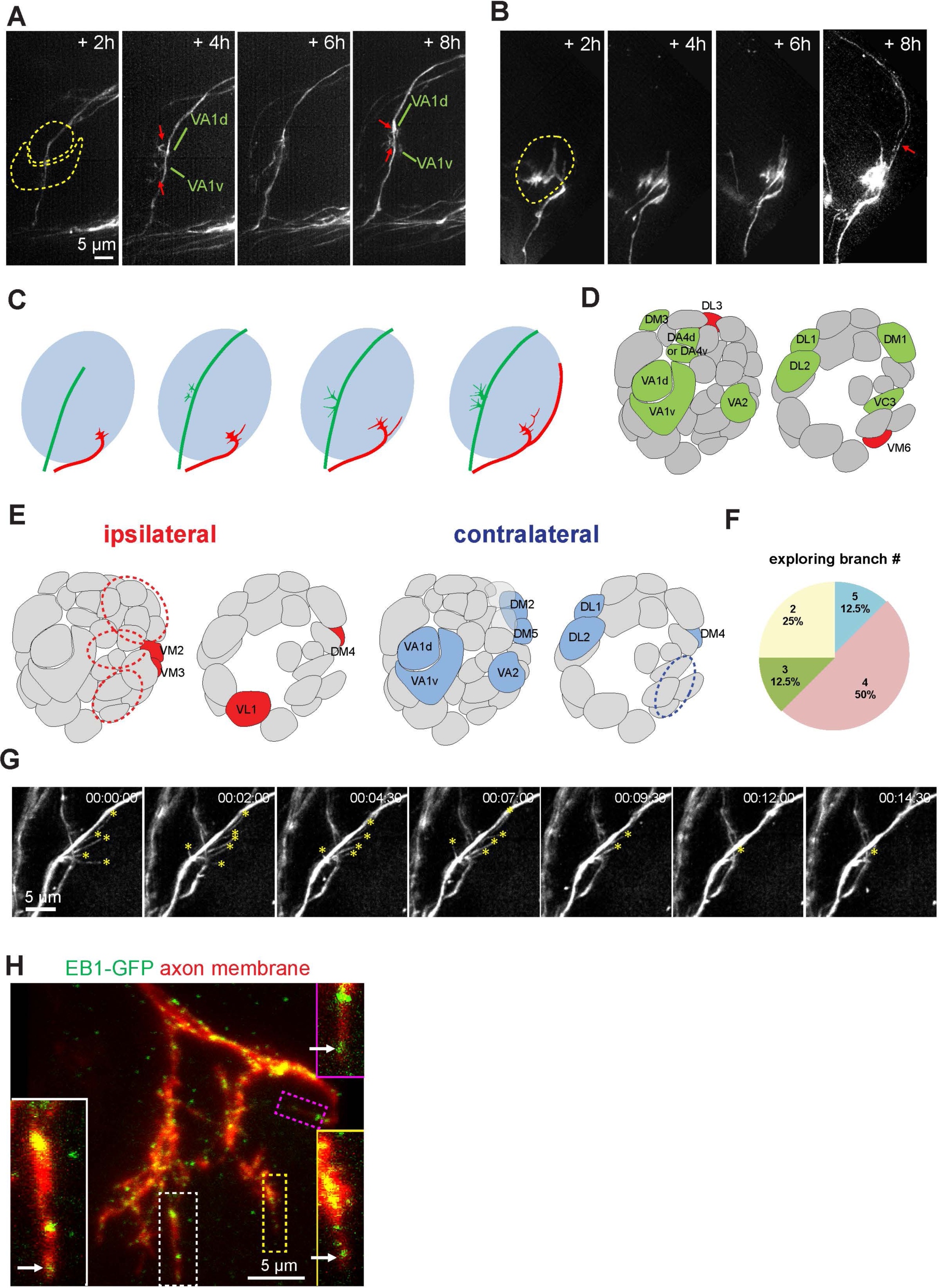
AO-LLSM imaging of ORN axon targeting, related to Figure 4, and confocal images of microtubule distribution, related to Figure 5. (A) Max intensity projection confocal images from 4 time points taken with AO-LLSM showing ipsilateral innervation of VA1d and VA1v ORNs. The main axon fascicles first pass the VA1d and VA1v glomeruli (dashed circles). Then two collateral branches innervate the glomeruli (arrows). (B) Max intensity projection confocal images from 4 time points taken with AO-LLSM showing ipsilateral innervation of a VM6 ORN. The axon first innervates the VM6 glomerulus (dashed circle). Then a collateral branch grows forward to cross the commissure (arrow). (C) Summary diagram of two collateral branching patterns: contralateral-projecting branch first (as in A, green axon) or ipsilateral branch first (as in B, red axon). (D) Summary map of ORN types that extend the contralateral projecting branch first (green) or ipsilateral innervation branch first (red) based on AO-LLSM imaging. One axon targeting to DA4d or DA4v cannot be determined due to the small size of the two glomeruli. (E) Summary map of ORN types captured for exploring branches in the ipsilateral or contralateral antennal lobe. Dashed ovals represent cases where the axons target a particular region of the antennal lobe but the target glomeruli were not identified. (F) Distribution of exploring branch number from single ORN axons. (G) Max intensity projection confocal images of a post-innervation terminus taken at indicated time points of a fast scanning session, showing gradual pruning of post-innervation branches. Asterisks mark the ends of post-innervation branches. (H) Max intensity projection confocal images of ORN axon terminals in fixed pupal brains from 34–36h APF labeled by *pebbled-GAL4*-based MARCM clones co-expressing membrane-Halo (red) and EB1- GFP, visualized with JF-Halo-646 ligand and anti-GFP antibodies. Arrows mark EB1-GFP puncta at the tips of post-innervation branches.

**Figure S7.**
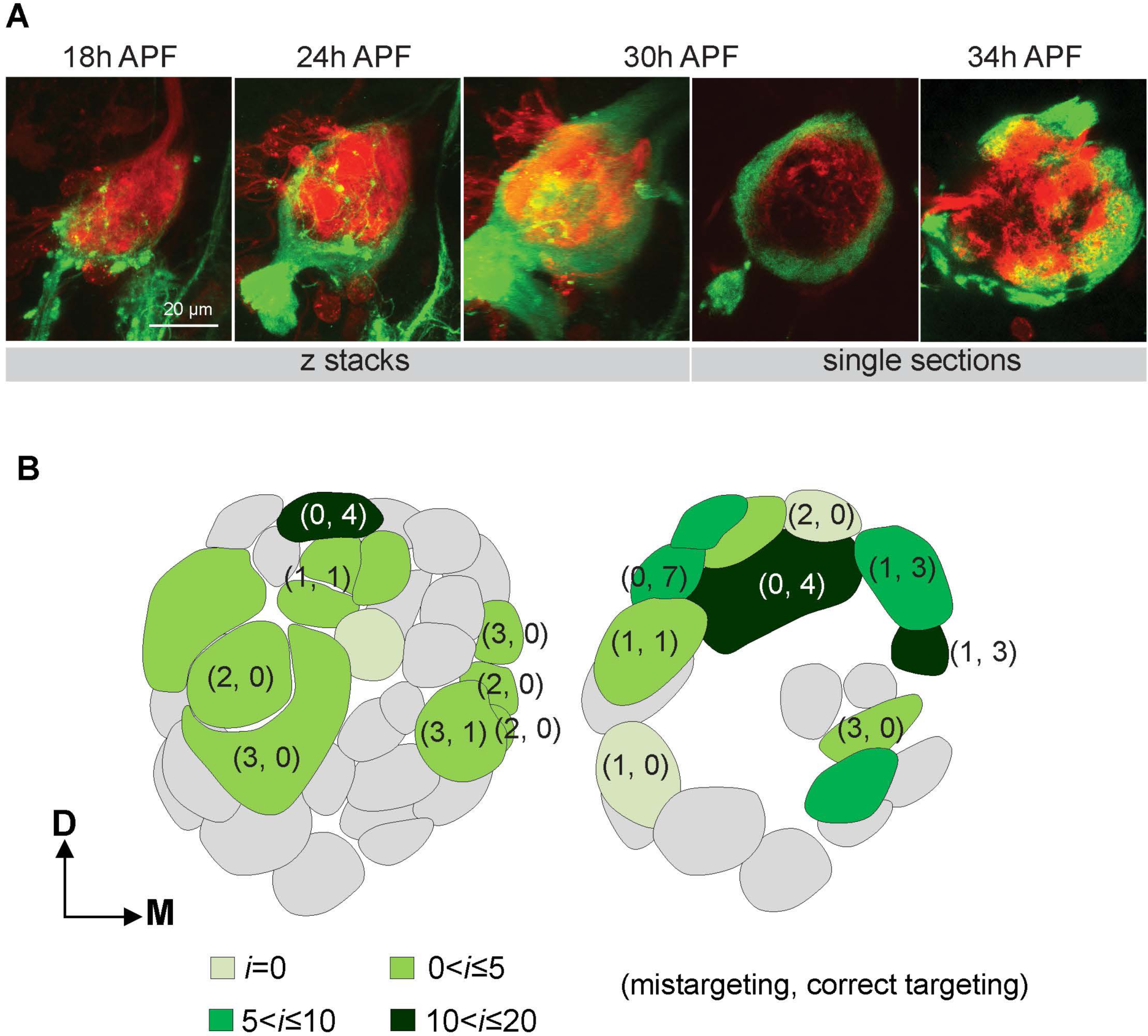
Time course of ORN axon development. (A) Projection of confocal images (left 3 images) or single optical section (right 2 images) from *pebbled-GAL4,UAS-mCD8-GFP* brains dissected at indicated time points, showing different developmental stages of ORN axons in the antennal lobe. GFP (green) and N-cadherin (red) were labeled. (B) Quantification of contralateral mistargeting and correct targeting of uncut ORN axons upon unilateral antennal nerve severing shown in an antennal lobe map with the ipsilateral branching waiting time *i* (Figure 3C) indicated with different green colors. First and second numbers in a bracket indicate the number of single axons that mistarget and target correctly from each type, respectively. All cases quantified were from explants with one antennal nerve severed at 30h APF followed by 24h culture ex vivo.

**Figure S8.**
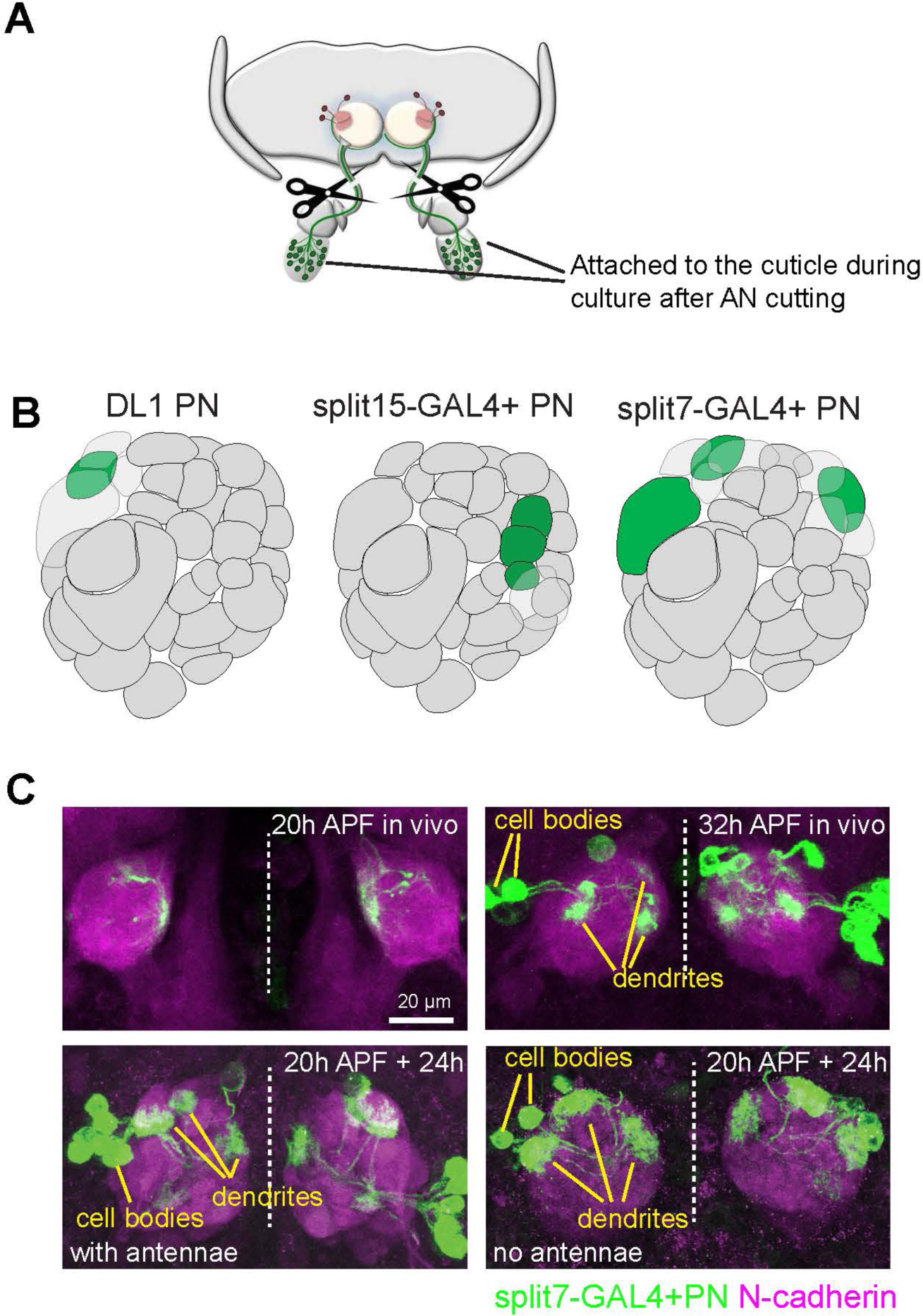
ORNs are not required for coarse targeting of PNs, related to Figure 7. (A) Schematic showing bilateral antennal nerve cutting. (B) Glomerular maps showing PN types labeled by single-cell MARCM clones (DL1) or by two split- GAL4s. (C) Max intensity projection confocal images of *split7-GAL4;UAS-mCD8-GFP* explants cultured for 24h with two antennae nerve intact or severed at the time of dissection at 20h. The bottom panels show *split7-GAL4;UAS-mCD8-GFP* brains dissected at 20h APF and 30h APF in vivo.

## Movie Legends

Movie 1. Time-lapse images showing targeting of *AM29-GAL4+* ORN axons during development. The first half of the movie shows that *AM29-GAL4+* ORN axons circumnavigate the antennal lobes and cross the midline. The second half of the movie shows that *AM29-GAL4+* ORN axons innervate DM6 and DL4 glomeruli. Dashed circles outline the antenna lobes and dashed vertical line marks the midline. Max intensity projections of image stacks taken every 20 min using a two-photon microscope. The two halves of the movie represent two different explants.

Movie 2. Time-lapse images showing targeting of a single DM2 ORN axon. Dashed circles outline the antennal lobes and dashed vertical line marks the midline. Max intensity projections of image stacks taken every 20 min using a two-photon microscope.

Movie 3. Time-lapse images showing targeting of a single DP1m ORN axon. Dashed circles outline the antennal lobes and dashed vertical line marks the midline. Asterisks denote transient collateral branches. Max intensity projections of image stacks taken every 20 min using a two-photon microscope.

Movie 4. Fast dynamics of the exploring branches during target searching. The first segment shows exploring branches from two ORN axons (same as Figure 4C; the right-most axon is unrelated). The second segment shows exploring branch from a single ORN axon in a different explant (same as Figure 4D). Dashed lines outline the shape of the antennal lobe. Max intensity projection image stacks taken every 30 sec using AO-LLSM.

Movie 5. Time-lapse images showing dendritic targeting of a single DL1 PN. The first half of the movie shows the active exploration of DL1 dendrites after initial innervation of the antennal lobe. The second half of the movie shows the decreased dynamics of DL1 dendrites during glomeruli boundary formation. Dendritic terminals and PN cell bodies were marked at the beginning of the movie. Max intensity projections of image stacks taken every 20 min using a two-photon microscope. The two halves of the movie show two different explants.

